# Organization of Areal Connectivity in the Monkey Frontoparietal Network

**DOI:** 10.1101/2020.06.30.178244

**Authors:** Bryan D. Conklin

**Affiliations:** Center for Complex Systems & Brain Sciences, Florida Atlantic University, Boca Raton, FL

**Keywords:** brain network, connectivity, cognitive neuroscience, macaque, connectome

## Abstract

Anatomical connectivity between cortical areas condition the set of observable functional activity in a neural network. The large-scale cortical monkey frontoparietal network (FPN) has been shown to facilitate complex cognitive functions. However, the organization of anatomical connectivity between areas in the FPN supporting such function is unknown. Here, a new connectivity matrix is proposed which shows the FPN utilizes a small-world architecture with an over-reliance on the M9 dynamical relay 3-node motif and degree distributions which can be characterized as single scale. The FPN uses its small-world architecture to achieve the kind of simultaneous integration and specialization of function which cognitive functions like attention and working memory require. Contrary to many real-world networks, the in and out single scale degree distributions illustrate the relatively homogeneous connectivity of each area in the FPN, suggesting an absence of hubs. Crucially, the M9 dynamical relay motif is the optimal arrangement for previously reported near-zero and non-zero phase synchrony to propagate through the network, serving as a candidate topological mechanism. These results signify the impact of the organization of anatomical connectivity in the FPN. They can serve as a benchmark to be used in the network-level treatment of neurological disorders where the types of cognition the FPN supports are impaired. Additionally, they can inform future neuromorphic circuit designs which aim to perform aspects of cognition.

## Introduction

The mammalian brain engages cortical neural networks during behavior^1,2^. The networks are composed of brain areas, or nodes, connected via axonal projections, or edges^3,4^. This structural organization of the network permits a set of functional interactions observed in neural signals recorded during behavior^5^.

A prominent functional interaction observed in neural signals is that of synchronous activity^6^. Recent studies have explored the structure-function relationship as it relates to synchrony through theoretical computational modeling. Vicente, Gollo, Mirasso, Fischer, & Pipa (2008) established that an apex node reciprocally connected to two unconnected nodes could foster zero-lag synchrony via dynamical relaying between the two unconnected nodes despite axonal conduction delays. This provided a topological mechanism for previously published findings from multicellular electrophysiological recordings which found distributed synchronous discharge in different structures of the cortex, hippocampus and thalamus^8,9^. Gollo, Mirasso, Sporns, & Breakspear (2014) extended this line of work by showing that a just a single resonance pair, two reciprocally connected nodes, could foster zero-lag synchrony in 3-node motifs. Further, they showed that the dynamical relay M9 motif, which has two resonance pairs, was optimally structured to provide both zero and non-zero phase lag synchrony. Finally, they found that the synchrony initiated locally with a resonance pair could propagate through the entire network, thereby impacting global network dynamics. These studies provide candidate topological mechanisms for the neural synchrony that has been reported to support cognitive functioning.

It is thought that an impaired structure-function relationship results in, or contributes to, various cognitive impairments. This phenomenon has been explored in aging, schizophrenia and autism^11–15^. A better understanding of the connectivity patterns that arise in mammals without mental health disorders can allow for comparison with the patterns which characterize impairments observed in disorders within the context of connectomics^16,17^.

Cognitive processing can occur through neural interactions both within and between regions of the cortex, forming large-scale cortical networks ^2,18,19^. The frontoparietal network (FPN) is one-such large-scale network comprised of sub-networks characterized by functional oscillatory dynamics that support aspects of cognition such as attention, cognitive control and working memory in both humans and non-human primates^20–25^. These dynamics exhibit unique spectral power and near-zero^26–28^ and non-zero^29^ phase-lag synchrony properties^24,30–34^. However, it is not understood how the unique structure of the FPN enables these properties to occur in the patterns required to support cognitive processing. The topological properties of this network remain to be elucidated.

Here the anatomical connections of the FPN are identified based on collated tract-tracing studies and examine the ways in which the topology acts as a reliable, integrative substrate while contributing to the reported neuronal dynamics which support varied cognitive functions. A new association matrix is proposed that uses a more finely grained parcellation scheme than previous studies^35^ with enough nodes for future analyses which require a high level of resolution^36^. A graph theoretic topological analysis^4,37^ was conducted to discover connectivity patterns in the 399 connections that make up the FPN and how they support cognitive functioning. The FPN is shown to be made up of relatively homogeneous connectivity between areas with an apparent lack of hub nodes controlling information flow. Further, the FPN utilizes a structural motif known for optimally promoting near-zero and non-zero neural synchrony. Finally, the FPN is discovered to be a small-world network, conferring both functional specialization and topological integration. Therefore, the FPN leverages a distributed connectivity architecture useful for critical information processing. It is optimally structured to support various aspects of cognition through neural synchrony and integration into coherent streams which support overall behavior.

## Results

### The Frontoparietal Connectome

To identify the structural connectivity of the FPN, axonal projections between and within the frontal and parietal regions of the monkey were collated using the results of tract-tracing studies on non-human primates (Methods; Supplementary Table 1) according to the parcellation scheme established by Petrides & Pandya (2007) (Figure 1).

**Figure 1.**
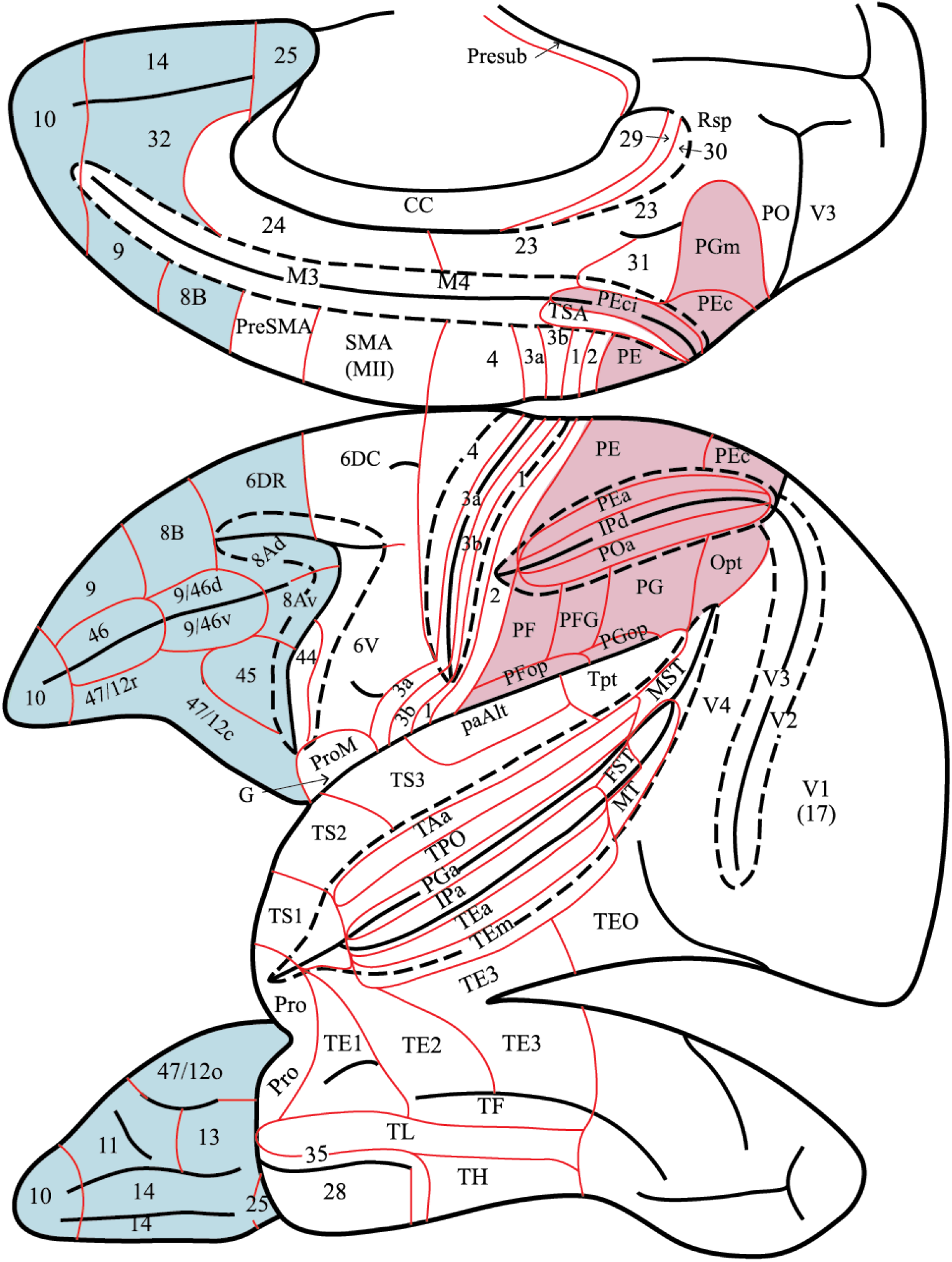
Lateral, orbital, and medial views of macaque cortex. Areas delineated by architectonic characteristics identified for the prefrontal cortex by Petrides & Pandya (1994), the posterior parietal cortex by Pandya & Seltzer (1982), the superior temporal gyrus by Pandya & Sanides (1973), the inferotemporal and superior temporal sulcus cortex by Seltzer & Pandya (1978), and the posterior parahippocampal gyrus by Rosene & Pandya (1983). The 17 frontal areas and 13 parietal areas that make up the frontoparietal network under examination in this study are colored blue and pink, respectively. *Adapted from Petrides & Pandya (2007) with permission*.

These connections were assembled into a binary, directed adjacency matrix comprised of 30 nodes, 17 frontal and 13 parietal (Figure 2).

**Figure 2.**
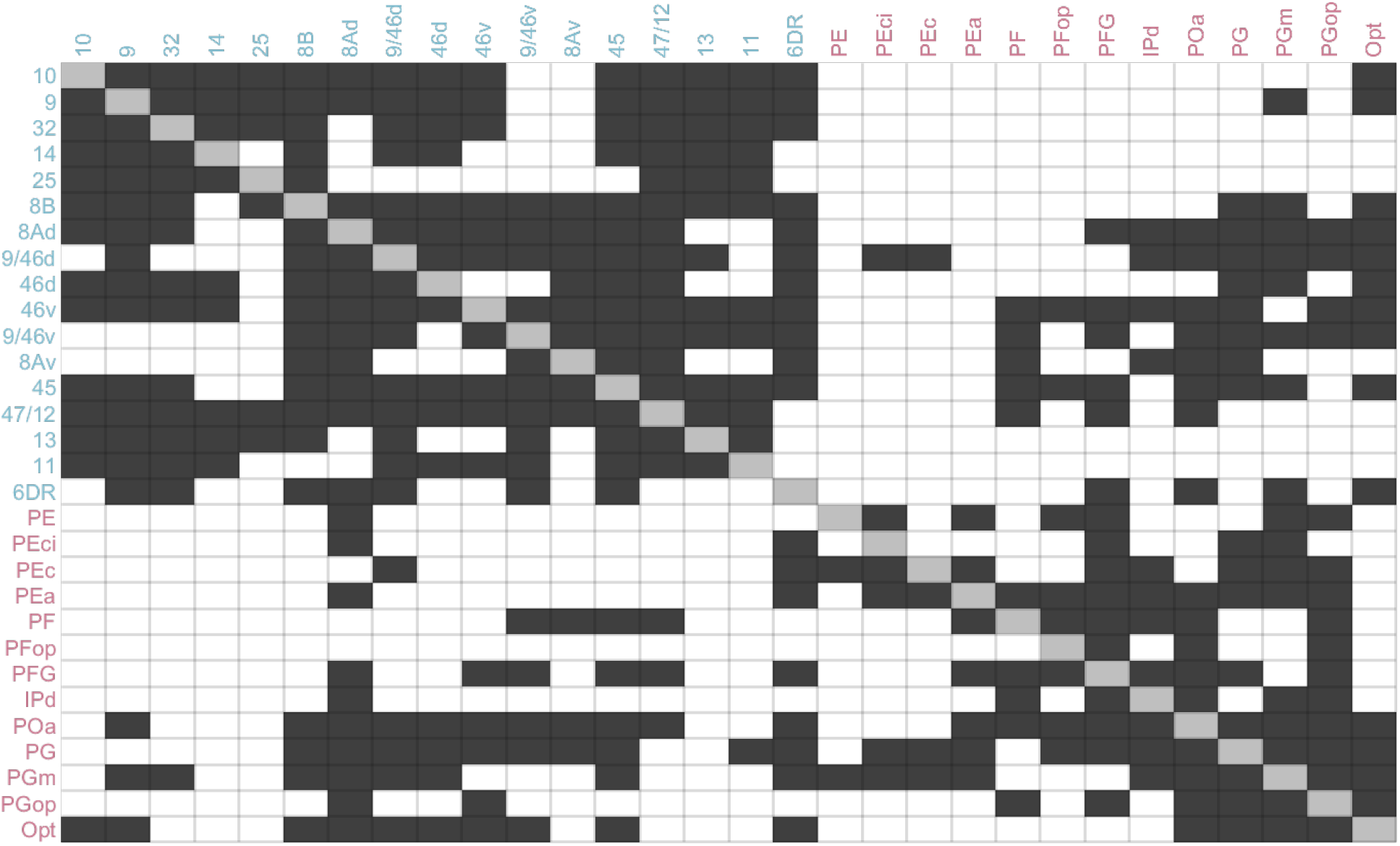
Binary, directed adjacency matrix describing the connectivity of the frontoparietal monkey network. The rows of the matrix represent source nodes, and the columns represent projection target nodes. All nodes represent architectonic areas from the Petrides & Pandya (2007) parcellation scheme. Connections are indicated with a black square. White squares represent connections that were not reported in the collated studies. Grey squares on the diagonal represent within-area connections and are not considered in the graph analysis. The 17 frontal areas are in blue, while the 13 parietal areas are in pink.

The adjacency matrix can be used to visualize network topology as a graph to better understand connectivity patterns (Figure 3). Here, nodes are sized according to their total degree, with bigger nodes representing areas with a greater number of incoming and outgoing connections than smaller nodes. Apart from a few parietal areas and one frontal, most of the areas in the FPN are of a similar size, representing similar total connectivity.

**Figure 3.**
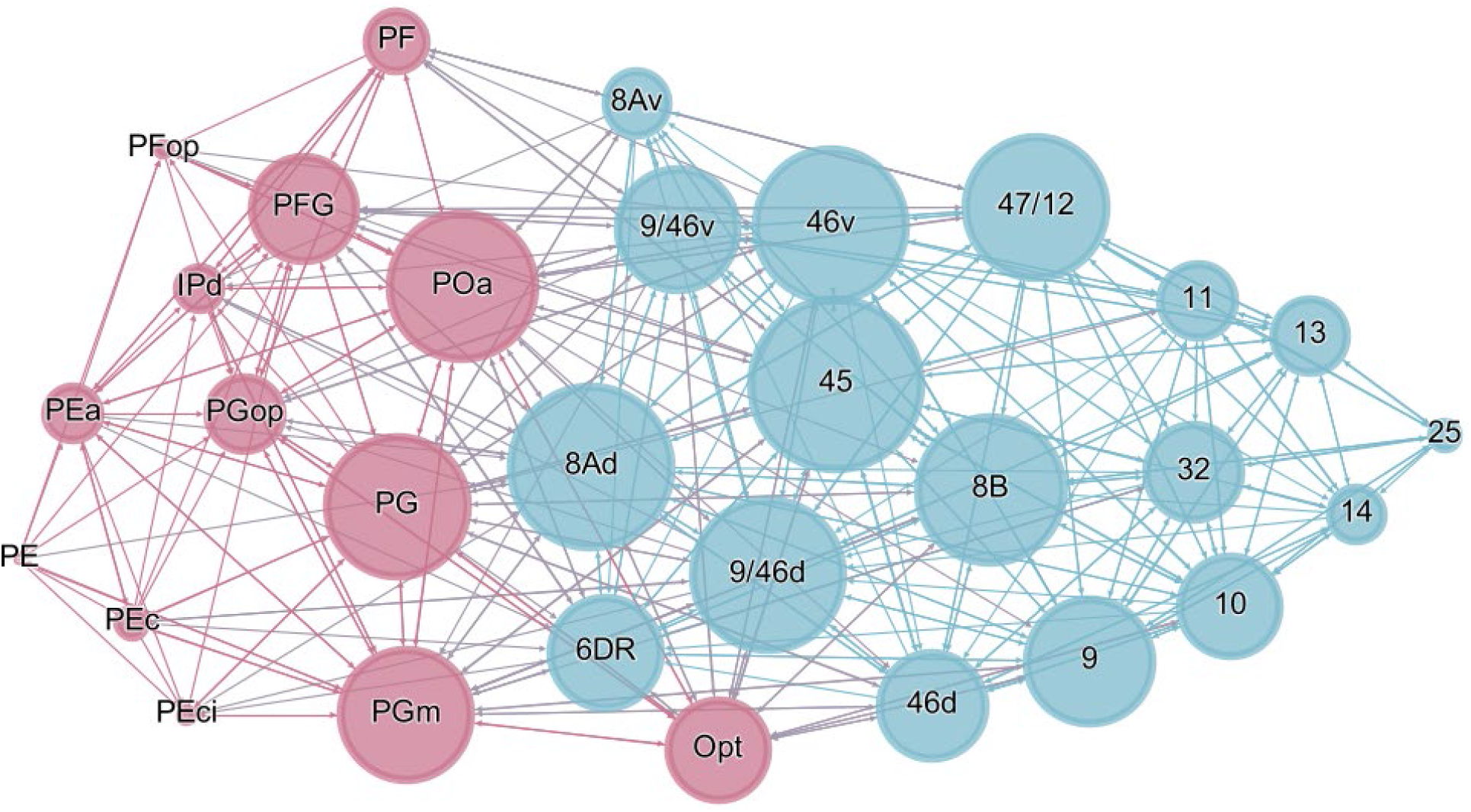
A graph representation of the frontoparietal network generated from the adjacency matrix. Frontal nodes are colored in blue and parietal in pink. The 399 directed edges between nodes represent projections between sources and targets. Node size is determined by total degrees, which is the sum of in-degree and out-degree connectivity. Bigger nodes represent a greater total degree than smaller nodes. The spatial distribution of the nodes is a function of the ForceAtlas layout in Gephi, a network visualization and analysis open-source software tool (Methods) The layout controls clustering and dispersion of nodes via attraction and repulsion strength parameter values^44^. Most nodes are of a similar size, representing relatively homogeneous connectivity.

### Graph Theoretic Analysis

Degree distributions convey information about how connectivity is allocated across the network^45^. The FPN’s in and out degree distributions can be characterized as either single-scale, scale-free or broad-scale. Single-scale distributions are highly unlikely to contain hubs, while scale-free and broad-scale distributions have a high likelihood of hub nodes^46^.

The topology of some brain networks^47–49^ present with hub nodes which serve to coordinate the majority of information transfer in the network^4^. These scale-free degree distributions are fit by a power law^46^ with no node exhibiting connectivity typical of other nodes. However, other studies have reported networks of the brain which do not follow power laws^50–52^.

Both tails of the degree distributions of the FPN were tested according to the Clauset, Shalizi, & Newman (2009) recipe to determine whether they were fit by a power law (Methods)^53^. The following parameter values were calculated for the in-degree distribution fit:
*α*= 3.375, *x*_*min*_ = 12 and *L* = −62.197 and for the out-degree distribution fit: *α*= 3.2521, *x*_*min*_ = 10 and *L* = −70.3376.

The degree distributions were visualized by plotting their complementary cumulative distribution functions (cCDF) on logarithmic axes^45^ with their estimated power-law fits^53^ (Figures 5a & b). The cCDF conveys the probability of finding a node with degree larger than some random value, *x*. Scale-free networks will approximate a straight line in these plots^54^.

**Figure 4.**
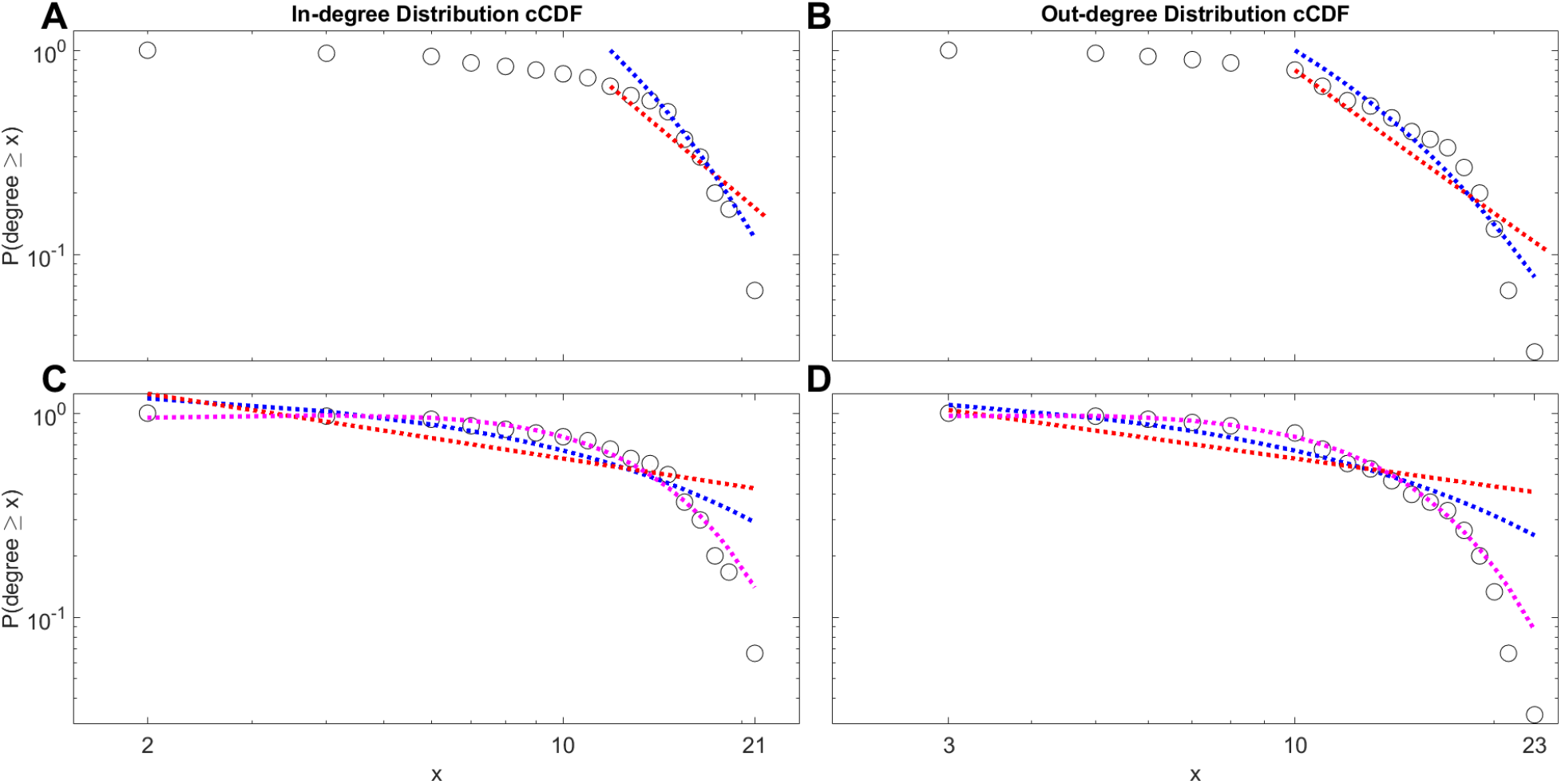
Complementary cumulative distribution function (cCDF) of the in-degree distribution (**A** & **C**) and out-degree distribution (**B** & **D**) for the FPN plotted on a logarithmic scale. Tail fits calculated according to the recipe proposed by Clauset et al., (2009) are shown in **A** & **B** and fits over the entire distribution are shown in **C** & **D**. Black circles represent the empirical distribution, the red dotted line represents a power-law fit, blue dotted line represents an exponential fit and the purple dotted line represents a Gaussian fit (**C** & **D** only). **A.** The tail fit begins from a lower bound estimate of x=12 (x_min_). However, it is not likely that the in-degree distribution follows a power law (*D* = 0.2097, *p* = 0.0158)^53^. Rather, a log-likelihood ratio shows an exponential distribution is a better fit to the tail of the distribution (*R* = −3.386, *p* = 0.00071)^55^. **B.** The tail fit begins from a lower bound estimate of x=10 (x_min_). The goodness-of-fit test does not rule out the possibility that the out-degree data may have been drawn from a power-law distribution (*D* = 0.1446, *p* = 0.1728)^53^. However, a log-likelihood ratio shows that an exponential fit may be just as good, or bad (*R* = −1.676, *p* = 0.094)^55^. **C.** A Gaussian model (adj R^2^ = 0.9740), fits the in-degree empirical data better than an exponential (adj R^2^ = 0.8293) or power law (adj R^2^ = 0.5660). **D.** A Gaussian model (adj R^2^ = 0.9909), fits the out-degree empirical data better than an exponential (adj R^2^ = 0.8981) or power law (adj R^2^ = 0.0.6988).

Next, the Kolmogorov-Smirnov statistic was used to test the goodness-of-fit between the empirical data and that drawn from a power-law distribution^53^. The in-degree data is not likely to have been drawn from a power-law distribution (*D* = 0.2097, *p* = 0.0158). However, a power-law distribution cannot be ruled out for the out-degree data (*D* = 0.1446, *p* = 0.1728). The data needs to be compared with other distributions to determine whether they match the data as well, or better.

Finally, different distributions were tested to see whether they were better fits to the tail of each distribution^55^. The log-likelihood ratio was calculated between the two candidate distributions. It was positive if the data was more likely in the first distribution and negative if more likely in the second distribution. Using this ratio, the exponential distribution was found to fit the tail of the in-degree data significantly better than a power-law distribution (*R* = −3.386, *p* = 0.00071). However, this does not mean the exponential distribution is an objectively good fit for the data. The ratio showed no significant difference between a power-law fit of the tail of the in-degree data and the following distributions: log-normal (*R* = −1.6299, *p* = 0.103), log-normal positive (*R* = −4.40, *p* = 0.1031) and truncated power-law (*R* = 3.693, *p* = 0.082). Despite the support for a power-law fit of the tail of the out-degree distribution based on the goodness-of-fit test in the previous step, the ratio showed the following distributions were no better or worse: exponential (*R* = −1.676, *p* = 0.094), log-normal (*R* = −0.978, *p* = 0.328), log-normal positive (*R* = −1.798, *p* = 0.328) and truncated power-law (*R* = −1.918, *p* = 0.083). It may be that all these distributions describe the data equally poorly.

Due to the poor fits to the candidate distributions using the tail-fitting recipe proposed by Clauset, Shalizi, & Newman (2009), the entire empirical in-degree (Figure 5c) and out-degree (Figure 5d) distribution data was also fit to power, exponential and gaussian models (Methods). These distributions qualify as potential fits based on their success in real-world connection distribution data^52,54^. The in-degree data was most consistent with a Gaussian distribution (adj R^2^ = 0.9740), in contrast to the exponential (adj R^2^ = 0.8293) and power law (adj R^2^ = 0.5660) distributions. The out-degree data was also most consistent with a Gaussian distribution (adj R^2^ = 0.9909), in contrast to the exponential (adj R^2^ = 0.8981) and power law (adj R^2^ = 0.6988) distributions.

### Motifs

Most studies analyze subgraphs of the larger monkey FPN due to technological limitations^24,30,32,56^. Some subgraphs may carry more importance in establishing the topological organization of the larger FPN than others. These subgraphs are known as structural motifs that serve as essential building blocks for the larger system^57,58^. They enable a variety of transient connection dynamics known as functional motifs^45^. Motifs of size 3 (M=3) are typically studied^58^. They are the most computationally tractable. The studies of the monkey FPN usually have at least 3 recording sites spanning the network. Additionally, the modeling work detailing the structure-function relationship through neural synchrony was based on size 3 motifs^7,10^. Accordingly, the FPN was analyzed to determine which structural motifs of size 3 were overrepresented in comparison with benchmark null networks. Any overrepresented motifs serve as the anatomical building block(s) of the network and give rise to specific functional interactions which can be observed in the literature. Further, regional areas that participate in these essential structural motifs can be important recording targets in electrophysiological studies that aim to describe cognitive behavior supported by the FPN.

To discover the anatomical building blocks used to create the FPN, structural motifs comprised of 3 areas or nodes were analyzed to determine which occurred with a significantly greater frequency than would be expected by chance. There are 13 possible motif classes of size three^58^.

As reported in other studies of cortical connectivity^58–60^, motif class ID 9 (M9) was found to be significantly overrepresented (*p* = 0, *z* = 12.0322 random networks, *p* = 0.0489, *z* = 1.6121 lattice networks) (Supplementary Table 2) in the empirical FPN (Figure 6). It forms an open triangle where two nodes are bidirectionally connected to a third apical node but are not connected to each other. The M9 motif is known as the dynamical relaying motif due to the unique functional capabilities it provides^7,61^. The M13 motif also appeared to be overrepresented, but not to a statistically significant extent (*p* = 0, *z* = 19.1224 random networks, *p* = 0.0538, *z* = 1.6108 lattice networks) (Supplementary Table 2).

**Figure 5.**
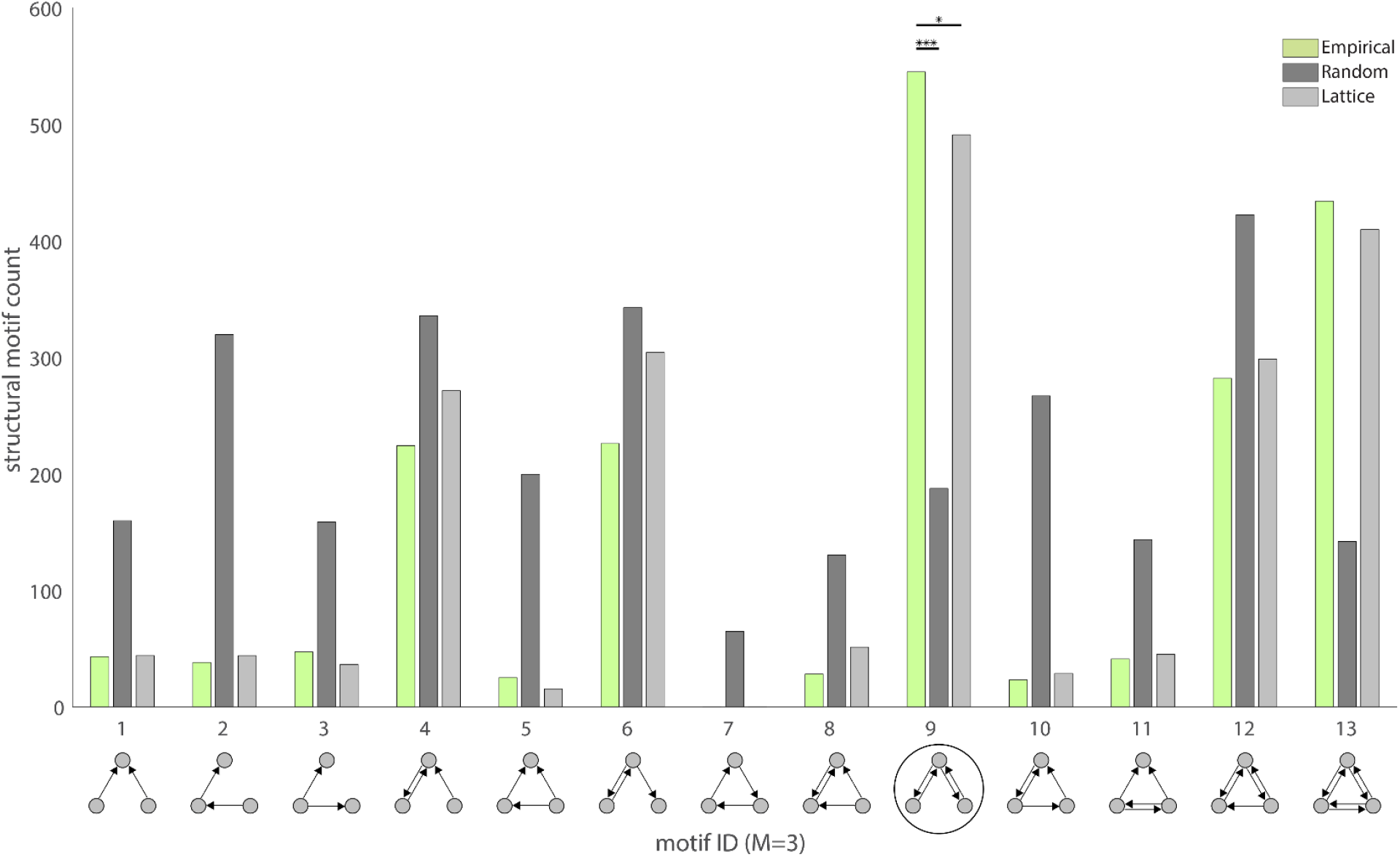
Comparison of the structural motif frequency spectrum for the empirical FPN with random and lattice benchmark networks. Motif class ID 9 was overrepresented in comparison to 100,000 random benchmark networks (*p* = 0, *z* = 12.032) and lattice networks (*p* = 0.049, *z* = 1.612). This dynamical relaying motif serves as a fundamental anatomical building block of the network, conferring a repertoire of functional dynamics such as neural synchrony that can be used to support various aspects of cognition. Empirical structural motifs counts are in green. Motif counts for random networks are in dark grey and the counts for lattice networks are in light grey.

### Small world

Networks demonstrating topology simultaneously consistent with a small characteristic path length and a high degree of clustering are called “small-world” networks ^62^. These networks cluster nodes into modules thereby minimizing wiring costs and efficiently integrating the topology. The result is an optimal information processing paradigm marked by fast transmission speeds and successful integration. There have been reports of many networks in neuroscience qualifying as small-world, such as the neural network of *C. elegans*, the brain-stem reticular formation and the mouse connectome ^52,63^.

Two indicators of “small-worldness” were calculated for the FPN using 100,000 null networks. First, Humphries’ index, σ (Methods), was estimated as 1.1429. A value greater than 1 is considered an indication that the network is small-world. Second, Telesford’s alternative index, w, was estimated as 0.0297. A value close to 0, specifically one that falls within the range of −0.5 ≤ ω ≤ 0.5, is considered an indication that the network is small-world. Negative values indicate a graph with more lattice-like characteristics and positive values indicate more random characteristics. Both metrics provide strong support for classifying the FPN as a small-world network with Telesford’s index showing a very slight skew towards random characteristics.

## Discussion

The tail of the in-degree distribution of the FPN does not appear to follow a power-law^53^, which can only occur in the presence of hub nodes. Additionally, because it is more likely to have been drawn from an exponential distribution, it is not considered heavy-tailed^55,64^. Heavy tailed distributions have many outliers. Exponential distributions are not likely to have outliers. Exponential decay in the tail of the degree distribution of neural networks has also been reported elsewhere^51,54^. It is important to keep in mind that this result only states the tail of the data is better fit by an exponential distribution than a power law. It does not imply that the exponential distribution is an objectively good fit.

The tail of the out-degree distribution of the FPN had moderate support for a power law fit. However, it was not significantly different from an exponential, log-normal, log-normal positive or truncated power-law fit either. These distributions may all be bad fits. At minimum, the results support the view that neither degree distribution follows a power law.

The entire in and out degree distributions yielded strong results for both a Gaussian and exponential fit^65^. The Gaussian fit may be stronger because it uses an extra parameter.

Notably, with both procedures yielding either Gaussian or exponential fits for the data, it is highly likely both distributions have a single, characteristic scale^54^. Single-scale systems have a very low probability of large deviations from a characteristic modal value, resulting in a relatively homogenous distribution of connectivity across nodes^45^.

Thus, the FPN is organized in such a way that the total number of incoming and outgoing connections to and from each of its nodes is relatively consistent across areas. There are only a small number of areas that greatly differ in the number of their total incoming and outgoing connections. Strategically, this could mean that information is diffused throughout the network, enabling concurrent or redundant processing. This creates a reliable substrate over which neural representations can be formed, maintained, and communicated^66^.

Conversely, scale-free networks with hub nodes contribute to a heterogenous connectivity profile. Simulations have shown that scale-free networks are more resilient against random node failures than single-scale networks^37,67^. This is because there are a much greater number of low-degree nodes in the scale-free networks leading to a greater likelihood that the random failures occur in these nodes. Therefore, there is little to no impact on overall network functionality. A disease that successfully impairs cognitive function supported by the single-scale FPN may owe its effectiveness to a random attack strategy.

The simulations also show that scale-free networks are much more vulnerable to targeted attacks on their highly connected hub nodes than the single-scale networks. The hub nodes represent single points of failure in the network. It is possible that serious diseases have evolved to target hub nodes, resulting in important cell death. This behavior may be what enables rapid disease progression in some patients. In a single-scale topology, if disease or other form of neural insult negatively impacts a single area, other similarly connected areas can pick up the slack, salvaging the integrity of the network and thereby supporting appropriate function.

The M9 dynamical relaying motif was statistically overrepresented in the FPN, establishing its importance as an anatomical building block of the network. Previous studies have established that its structure is ideal for providing both zero-lag synchrony between its driven nodes and non-zero phase synchrony between its relay and either driven node^7,10^.

The synchrony initiated by M9’s resonance pair can extend to nodes upwards of four steps removed with very little synchrony decay^10^. Therefore, it may serve as the mechanism for zero and non-zero lag synchrony throughout the entire FPN. Crucially, this establishes the M9 motif as a proposed generative topology which likely gives rise to the synchrony previously reported in subgraphs of the FPN supporting attention^23^, categorization^32^, working memory^24,31^ and cognitive control^68^.

Finally, of the 3-node motifs with resonance pairs, the M9 class was observed to be the most robust to large conduction delays and delay mismatches^10^, which are a defining feature of the long-range cortical FPN.

Overall, the M9 motif is a maximally versatile building block for the FPN, conferring global near-zero and non-zero synchrony, enabling robust streams of communication and binding which support multiple aspects of cognition.

The FPN’s small-world architecture provides an economic^69^ substrate for the brain to process information while minimizing wiring cost. Information can flow efficiently due to a low global path length and integrate in functionally specialized modules due to high clustering. This serves as an elegant solution to the problem of integrating function while allowing for concurrent specialization. The simultaneous specialization and integration of function has been observed empirically in studies of cognition in both humans and non-human primates^23,70^.

Notably, despite a relatively homogeneous degree distribution with no clear evidence of hub nodes, there was enough dispersed integration to qualify the network as small world. The Telesford index did identify slightly random characteristics for the network, which supports the degree distribution findings. Ultimately, this suggests that small-world networks do not need to have nodes with much larger degrees relative to the rest of the network. If the nodes all have a sufficiently large relative number of connections, this provides the necessary integration to qualify as small-world.

Finally, small-world characteristics have been shown to better support oscillatory synchronization in contrast to random or lattice networks^71,72^. Thus, in addition to the synchrony advantage afforded by the over-represented M9 motif, the small-world qualification permits the FPN to optimally conduct neural synchrony in support of cognition.

In conclusion, the monkey FPN is uniquely anatomically constructed to support various important cognitive functions. Its fundamental building block is a dynamical relaying motif which confers the optimal 3-node connectivity profile for the synchrony necessary to support types of cognition like working memory and attention. Its small-world architecture provides the integration and specialization of function that complex cognitive tasks require. Finally, the degree of connectivity in the network is dispersed, establishing a reliable substrate for engaging in complex cognitive tasks.

Understanding the structural mechanism which gives rise to the functions a network is known to support provides a pathway to examine various neurological disorders. In diseases such as schizophrenia, Parkinson’s disease or Alzheimer’s disease, aspects of cognition that the FPN is known to support such as working memory and attention are impaired. Using the results of this study, scientists can examine the ways in which the organization of anatomical connectivity between areas in the FPN are different in pathological animal models or patients suffering from these disorders. Treatments could be devised that impact the pathological network in order to bring it closer to exhibiting the properties that networks in healthy individuals show^73^.

Additionally, engineers interested in exploring the mechanisms of cognition at a network level can use these results to explore their physical instantiation in machines. It would be interesting to see how the FPN may be realized in a neuromorphic circuit which emphasizes homogeneous connectivity in a small-world architecture using a dynamical relaying motif as its core building block^74^.

Finally, future empirical studies may want to use the FPN connectivity matrix presented here to identify subgraph regions of interest for recording during cognitive tasks. For instance, a research team interested in studying working memory neural dynamics may want to target areas that form the M9 motif so that the origination of any near-zero or non-zero phase lag synchrony contributing to working memory function is more likely to be identified.

The author hopes that in the future this level of graph theoretic analysis will be applied to other large-scale cortical networks supporting complex functions.

## Methods

### Parcellation Scheme

Non-human primate parcellation schemes are numerous and varied. Some of the ways they are derived include establishing boundaries based on cytoarchitecture, myeloarchitecture, or function^35,75–81^. This study uses the parcellation scheme identified in Petrides & Pandya (2007) (Figure 1). In this scheme, there are sixteen prefrontal areas: 10, 9, 32, 14, 25, 8B, 8Ad, 9/46d, 46d, 46v, 9/46v, 8Av, 45, 47/12, 13, 11, one frontal area 6DR and thirteen parietal areas: PE, PEci, PEc, PEa, PF, PFop, PFG, IPd, POa, PG, PGm, PGop, Opt. The prefrontal areas were delineated based on architectonic characteristics identified by Petrides & Pandya (1994) and the posterior parietal areas based on Pandya & Seltzer (1982). Area 6DR was included for its hypothesized involvement in the FPN^24^.

This scheme was chosen because it allows for a level of granularity sufficient for exploring possible functional implications^82,83^ and it provides enough areas (30) for a graph theoretic analysis^84^. For instance, analyses such as the Clauset, Shalizi, & Newman (2009) recipe for analyzing power-law distributed data require enough nodes to ensure the accuracy of the technique^55^. Each delineated area is considered a node. The connections between them are called edges.

### Connections

Published tract-tracing studies were collated, and results were interpreted according to the Petrides & Pandya (2007) parcellation scheme. The studies utilized both retrograde and anterograde tracing techniques. Retrograde tracers injected into an area migrate back along axons to the neuronal somas they originate from^85^. Anterograde tracers injected into an area are transported away from the soma to their site of termination^86^. The resulting labeled neurons or boutons can then be seen under a microscope allowing for statements to be made about direct axonal projections between areas.

If a tracer clearly had uptake into adjacent areas, its results were not considered in the analysis. Tracer injection sites were defined according to the author’s interpretation of a mapping to the selected parcellation scheme. Many connections were present in multiple studies. However, others were only reported in a single study due to the small number of tract-tracing investigations on particular areas of cortex represented in the parcellation scheme (Supplementary Table 1).

It is important to keep in mind that most studies involved Old World macaque monkeys (Macaca fascicularis, Macaca mulatta, or Macaca nemestrina)^87–89^, but a small number also involved New World monkeys such as squirrel monkeys (Saimiri sciureus)^90^ or owl monkeys (Aotus trivirgatus)^91^. Both kinds are of the infraorder Simiiformes. Furthermore, the connections were collated from both hemispheres, across sexes and age groups. It was our intention to provide a broad overview of the connections comprising the non-human primate frontoparietal network irrespective of variables such as non-human primate species, sex, age, or hemisphere.

Existing resources for obtaining connectivity information on the FPN were analyzed for efficacy. First, the work presented in Markov et al. (2014) served as an excellent example of a comprehensive whole-brain network analysis and provided valuable connectivity information. However, there were not enough unique areas injected to be useful to examine the frontoparietal network, specifically.

Next, the Collation of Connectivity data for the Macaque (CoCoMac) database was considered for obtaining connectivity information^92,93^. This database serves as a repository for macaque neural connections reported in the tract-tracing literature. To deal with the problem of inconsistently named areas, boundary conflicts and differing resolutions from parcellation schemes chosen by researchers, CoCoMac uses a routine that attempts to automatically map connections between schemes. This gives CoCoMac the ability to make connection statements which apply to specific parcellation schemes. Unfortunately, the accuracy of these statements is questionable, making them unreliable for use in analysis. For instance, if a study using a parcellation scheme different from that proposed by Petrides & Pandya (2007) reports that area 46 projects to area LIP (lateral intraparietal sulcus), CoCoMac won’t be able to make a reliable statement about whether the connection should be from 46d, 46v, 9/46d or 9/46v to area POa, the area which corresponds to LIP^94^. Clearly, the original study in this example lacks the specificity necessary to report a connection which applies to the selected parcellation scheme. The only solution is to re-interpret the finding from the original study using the selected parcellation scheme. Accordingly, all tract-tracing studies were re-interpreted, as necessary.

Unfortunately, there is no universally accepted method for defining connection strength between areas, despite recent interesting approaches^35^. The studies used in this analysis reported binary or qualitative descriptions of axonal connectivity. This results in the unfortunate situation where a dense connection between two areas is treated the same as a sparse connection. The binary directed matrix that results from this scenario, however, is still more informative than its undirected version, which is primarily used in fMRI studies with diffusion tensor imaging. The binary directed matrix reports a connection as either existing (with a 1) or not discovered (with a 0). It is important to note that the absence of a connection does not mean there is no projection between the two areas. Rather, it means that a connection between the areas was not reported.

### Graph Visualizations

The spacing of the nodes in Figure 3 is controlled by the ForceAtlas layout in Gephi, an open-source network visualization and analysis software tool^95^. Force-directed layouts, like that used in the ForceAtlas layout, quantify levels of spatial attraction and repulsion for the nodes based on some measure of pair-wise node distance like topological path length, the minimum number of edges between any two nodes^45^. These layouts minimize edge crossings, enhance symmetry, keep edge lengths uniform, spatially distribute nodes in a uniform manner and correlate node position with their topological adjacency.

### Graph Measures

All graph theoretic analyses were conducted using the publicly available Brain Connectivity Toolbox^96^.

### Degree Distribution

To determine whether the FPN’s degree distributions were scale-free or broad-scale, an analysis was conducted to test whether its in or out-degree distributions follow a power law. The recipe for evaluating the existence of power-law scaling proposed by Clauset, Shalizi, & Newman (2009) was followed using the publicly available MATLAB and Python^55^ scripts. The recipe is to first fit a power-law to the data above some lower bound, *x*_*min*_, using maximum likelihood estimation, then test its goodness of fit and finally to compare the power law with alternative distributions using a likelihood ratio test. A lower bound is used because it is typical for empirical data to only follow a power law for values above some level of *x*^53^. This results in a fit for the tail of the distribution.

First, the data is fit to a power-law using maximum likelihood estimation using the publicly available MATLAB function ‘plfit’.

The fitting procedure estimates parameter values for the following equation:

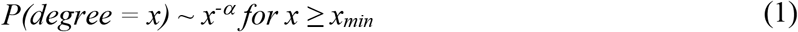

Where *α* is the maximum likelihood estimate of the scaling exponent, *x*_*min*_ is the estimated degree of the lower bound of power-law behavior and *L* is the log-likelihood (equation 3.5 in Clauset et al., 2009) of the data *x* ≥ *x*_*min*_ under the fitted power law.

Next, the goodness of fit between the empirical data and that drawn from a power-law distribution is tested using the Kolmogorov-Smirnov statistic in the publicly available MATLAB function ‘plpva’. There were 5,000 semiparametric repetitions of the fitting procedure. The statistic provides a p-value that allows a determination to be made about whether the power-law hypothesis can be ruled out. If *p* ≤ 0.1, the power law can be ruled out

Finally, the power law fit is compared with alternative distributions using a log-likelihood ratio test enabled via the Powerlaw Python package^55^. The distributions’ tails are compared to see which is a better fit to the data. If the log-likelihood ratio is positive, the data was more likely in the first distribution. If it is negative, the data was more likely in the second distribution. The p-value indicating directional significance is also provided.

Additionally, the entire in and out-degree distribution data were fit to power, exponential and gaussian models using MATLAB’s method of nonlinear least squares error minimization^65^, which minimizes the following:

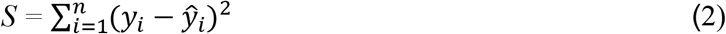

Where *n* is the number of empirical data points, y_i_ is the probability of a node’s degree being greater than or equal to a random degree (the cCDF), ŷ_i_ is the predicted probability value and *S* is the summed square of residuals.

### Null Models

The FPN was rewired using two different algorithms to generate null distributions used in the motif and small world analyses. Each algorithm creates networks at either end of the topological spectrum: random and lattice based^62^. The first algorithm functions by rewiring two random connections involving four nodes to a new connection scheme involving the same four nodes ensuring the connections in the new scheme do not already exist. If they do exist, the process is abandoned and begun anew with two different random connections. The algorithm proceeds in this fashion, maintaining the same network size, connection density and in and out-degree distributions as the original network, resulting in a random network^97^. It has been shown that completing the rewiring process at least 10 times the number of edges in the network is a long enough time to wait to ensure sufficient mixing^98^. The FPN has 399 edges. So, there were 3,990 iterations used to generate each null network.

The second algorithm functions in much the same way as the first. After rewiring the two random connections and ensuring they are novel, it must also be confirmed that they are now closer to the diagonal of the adjacency matrix than they were previously. This ensure connections only link nearby nodes. If this is not the case, the process is abandoned and begun anew with two different random connections. Just like the first algorithm, the network size, connection density and degree distributions are maintained. The result is a latticed network^50^.

### Motifs

All 13 classes of structural motifs of size three^58^ were tested for overrepresentation in the empirical FPN. The number of instances of each motif class were counted in the FPN, yielding a motif spectrum. These empirical frequencies were compared to 100,000 random and lattice benchmark networks’ motif spectra. P-values for each motif class were calculated as the fraction of times the frequency count of a benchmark network was higher than the count in the empirical FPN. If the p-value was less than 0.05 for both the random and lattice benchmark networks, the motif class was considered to be significantly overrepresented in the empirical FPN.

The amount of times each area participated across the 13 classes was quantified in the empirical FPN and compared to the frequency observed in the 100,000 random and lattice benchmark networks as well. P-values were calculated as the fraction of times the frequency count of each area’s participation in motif classes exceeded that observed in the empirical FPN. If the p-value was less than 0.05 for both the random and lattice benchmark networks, the area was considered to be significantly overrepresented in the empirical FPN.

### Small world

The small-world topology of a network can be quantified by comparing its observed characteristic path length, *L*, and clustering coefficient, *C*, with their distributions in null networks.

Average path length measures how efficiently information can be routed in a network. For any nodes *i* and *j*, the shortest path length, *l*_*i,j*_, between them is defined as the total number of edges one must traverse in navigating from node *i* to node *j* along the quickest route in the binary directed graph of the network. This can be computed for each node in a network algorithmically^99^. Next, each node’s shortest path length is averaged to quantify the average path length of the entire network, the characteristic or global path length:

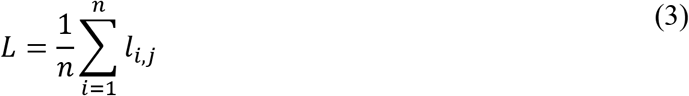

The clustering coefficient measures a network’s level of integration in terms of 3-node subgraphs. For any node *i*, its clustering coefficient is calculated using the following ratio:

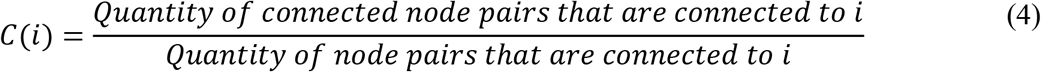

Next, each node’s clustering coefficient is averaged to quantify the clustering of the entire network:

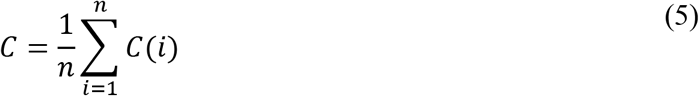

There are two metrics used to quantify the small-worldness of a network: Humphries’ index of small-worldness, σ^52^ and Telesford’s alternative index, ω^100^. The calculation of Humphries’ index requires the characteristic path length and clustering coefficient be normalized to their average values in appropriately randomized null networks. The following normalized values are obtained:

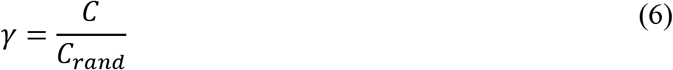

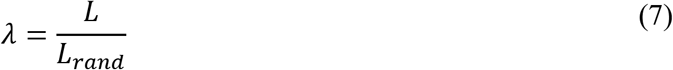

Where *C*_*rand*_ and *L*_*rand*_ are the average clustering coefficient and characteristic path length calculated for an ensemble of appropriately randomized null networks. Next, the normalized values are divided, providing the following index:

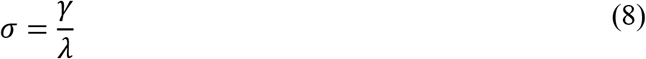

From this equation, if *λ* ~ 1 (low path length) and *γ* > 1 (high clustering), then *σ* > 1. Values of *σ* > 1 are typically considered an indicator of a network’s small-world organization.

The calculation of Telesford’s alternative index stems from the idea that appropriately matched lattice networks are better suited to normalize the clustering coefficient metric than randomized networks. This is because a lattice network presents with maximal clustering, while a random network presents with minimum global path length. Hence, Telesford et al. (2011) propose the following index:

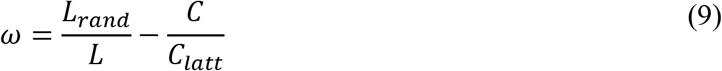

Where *C*_*latt*_ and *L*_*rand*_ are the average clustering coefficient and characteristic path length calculated for an ensemble of appropriately matched lattice and randomized null networks, respectively. The index ranges between −1 and 1. A value close to 0 is typically considered an indicator of a network’s small-world organization.

## Acknowledgements

Thank you William H. Alexander, Edward E. Ester and Randy D. Blakely for proofreading and providing helpful comments on the manuscript.

## Code Availability

No software was used to collect data for this study. However, software was used for the data analysis. The commercial product MATLAB R2019b was used for all analyses. The publicly available Brain Connectivity Toolbox v2019-03-03 was used for all graph theoretic analyses. The open-source Gephi v0.9.2 platform was used for graph visualizations. Aaron Clauset & Cosma Shalizi’s publicly available MATLAB functions v2012-01-17 and Jeff Alstott’s open-source powerlaw Python package v2020-02-02 were used for the degree distribution tail-fitting. All associated MATLAB code for this study can be found on the publicly available Github repo: https://github.com/thor4/Monkey-Data/tree/master/structural

## Data Availability

The data used in the analysis for this study can be found on the publicly available Github repo linked above. It can also be seen in Figure 1. Steps to collect the data were documented in the Methods section with corresponding supplementary tables.

## Author Contributions

BDC conceived of the study, collected the data, conducted the graph theoretic analysis and wrote the manuscript.

## Competing Financial Interests

There are no competing financial interests.

## Notes

https://github.com/thor4/Monkey-Data

### Competing Interest Statement

The authors have declared no competing interest.

## References

1. Buzsáki, G. & Draguhn, A. Neuronal oscillations in cortical networks. Science 304, 1926–9 (2004).

2. Bressler, S. L. & Menon, V. Large-scale brain networks in cognition: emerging methods and principles. Trends Cogn. Sci. 14, 277–290 (2010).

3. Sporns, O., Chialvo, D. R., Kaiser, M. & Hilgetag, C. C. Organization, development and function of complex brain networks. Trends Cogn. Sci. 8, 418–425 (2004).

4. Newman, M. Networks. (2018).

5. Wang, Z., Dai, Z., Gong, G., Zhou, C. & He, Y. Understanding structural-functional relationships in the human brain: A large-scale network perspective. Neuroscientist 21, 290–305 (2015).

6. Varela, F., Lachaux, J. P., Rodriguez, E. & Martinerie, J. The brainweb: Phase synchronization and large-scale integration. Nat. Rev. Neurosci. 2, 229–239 (2001).

7. Vicente, R., Gollo, L. L., Mirasso, C. R., Fischer, I. & Pipa, G. Dynamical relaying can yield zero time lag neuronal synchrony despite long conduction delays. Proc. Natl. Acad. Sci. U. S. A. 105, 17157–17162 (2008).

8. Traub, R. D., Whittington, M. A., Stanford, I. M. & Jefferys, J. G. R. A mechanism for generation of long-range synchronous fast oscillations in the cortex. Nature 383, 621–224 (1996).

9. Contreras, D., Destexhe, A., Sejnowski, T. J. & Steriade, M. Control of spatiotemporal coherence of a thalamic oscillation by corticothalamic feedback. Science (80-.). 274, 771–774 (1996).

10. Gollo, L. L., Mirasso, C., Sporns, O. & Breakspear, M. Mechanisms of Zero-Lag Synchronization in Cortical Motifs. PLoS Comput. Biol. 10, (2014).

11. Nakagawa, T. T., Jirsa, V. K., Spiegler, A., McIntosh, A. R. & Deco, G. Bottom up modeling of the connectome: Linking structure and function in the resting brain and their changes in aging. Neuroimage 80, 318–329 (2013).

12. Persson, J. et al. Structure-function correlates of cognitive decline in aging. Cereb. Cortex 16, 907–915 (2006).

13. Anderson, J. S. et al. Decreased interhemispheric functional connectivity in autism. Cereb. Cortex 21, 1134–1146 (2011).

14. Ben Bashat, D. et al. Accelerated maturation of white matter in young children with autism: A high b value DWI study. Neuroimage 37, 40–47 (2007).

15. Fornito, A., Zalesky, A., Pantelis, C. & Bullmore, E. T. Schizophrenia, neuroimaging and connectomics. NeuroImage 62, 2296–2314 (2012).

16. Reinhart, R. M. G. & Nguyen, J. A. Working memory revived in older adults by synchronizing rhythmic brain circuits. Nat. Neurosci. 22, (2019).

17. Polanía, R., Nitsche, M. A., Korman, C., Batsikadze, G. & Paulus, W. The importance of timing in segregated theta phase-coupling for cognitive performance. Curr. Biol. 22, 1314–1318 (2012).

18. Mesulam, M. Large-scale neurocognitive networks and distributed processing for attention, language, and memory. Ann. Neurol. 28, 597–613 (1990).

19. Mishkin, M. & Ungerleider, L. G. Contribution of striate inputs to the visuospatial functions of parieto-preoccipital cortex in monkeys. Behav. Brain Res. 6, 57–77 (1982).

20. Marek, S. & Dosenbach, N. U. F. The frontoparietal network: function, electrophysiology, and importance of individual precision mapping. Dialogues Clin. Neurosci. 20, 133–140 (2018).

21. Zanto, T. P. & Gazzaley, A. Fronto-parietal network: flexible hub of cognitive control. Trends Cogn. Sci. 17, 602–603 (2013).

22. Corbetta, M. Frontoparietal cortical networks for directing attention and the eye to visual locations: Identical, independent, or overlapping neural systems? Proc. Natl. Acad. Sci. U. S. A. 95, 831–838 (1998).

23. Fiebelkorn, I. C. & Kastner, S. A rhythmic theory of attention. Trends Cogn. Sci. 23, 1–36 (2019).

24. Salazar, R. F., Dotson, N. M., Bressler, S. L. & Gray, C. M. Content-Specific Fronto-Parietal Synchronization During Visual Working Memory. Science (80-.). 338, 1097–1100 (2012).

25. Lundqvist, M. et al. Gamma and Beta Bursts Underlie Working Memory. Neuron 90, 152–164 (2016).

26. Singer, W. & Gray, C. M. Visual feature integration and the temporal correlation hypothesis. Annu. Rev. Neurosci. 18, 555–586 (1995).

27. König, P., Engel, A. K. & Singer, W. Relation between oscillatory activity and long-range synchronization in cat visual cortex. Proc. Natl. Acad. Sci. U. S. A. 92, 290–294 (1995).

28. Singer, W. Neuronal Synchrony: A Versatile Code for the Definition of Relations? Neuron 24, 49–65 (1999).

29. Bastos, A. M., Vezoli, J. & Fries, P. Communication through coherence with inter-areal delays. Current Opinion in Neurobiology 31, (2015).

30. Kornblith, S., Buschman, T. J. & Miller, E. K. Stimulus load and oscillatory activity in higher cortex. Cereb. Cortex 26, 3772–3784 (2016).

31. Jacob, S. N., Hähnke, D. & Nieder, A. Structuring of Abstract Working Memory Content by Fronto-parietal Synchrony in Primate Cortex. Neuron 99, 588–597.e5 (2018).

32. Antzoulatos, E. G. & Miller, E. K. Synchronous beta rhythms of frontoparietal networks support only behaviorally relevant representations. Elife 5, 1–22 (2016).

33. Fries, P., Roelfsema, P. R., Engel, A. K., König, P. & Singer, W. Synchronization of oscillatory responses in visual cortex correlates with perception in interocular rivalry. Proc. Natl. Acad. Sci. U. S. A. 94, 12699–12704 (1997).

34. Fries, P., Womelsdorf, T., Oostenveld, R. & Desimone, R. The effects of visual stimulation and selective visual attention on rhythmic neuronal synchronization in macaque area V4. J. Neurosci. 28, 4823–4835 (2008).

35. Markov, N. T. et al. A weighted and directed interareal connectivity matrix for macaque cerebral cortex. Cereb. Cortex 24, 17–36 (2014).

36. Cavada, C. & Goldman-Rakic, P. S. Posterior parietal cortex in rhesus monkey: II. Evidence for segregated corticocortical networks linking sensory and limbic areas with the frontal lobe. J. Comp. Neurol. 287, 422–445 (1989).

37. Albert, R. & Barabási, A. L. Statistical mechanics of complex networks. Rev. Mod. Phys. 74, 47–97 (2002).

38. Petrides, M. & Pandya, D. N. Efferent Association Pathways from the Rostral Prefrontal Cortex in the Macaque Monkey. J. Neurosci. 27, 11573–11586 (2007).

39. Petrides, M. & Pandya, D. N. Comparative architectonic analysis of the human and the macaque frontal cortex. in Handbook of neuropsychology (1994).

40. Pandya, D. N. & Seltzer, B. Intrinsic connections and architectonics of posterior parietal cortex in the rhesus monkey. J. Comp. Neurol. 204, 196–210 (1982).

41. Pandya, D. N. & Sanides, F. Architectonic parcellation of the temporal operculum in rhesus monkey and its projection pattern. Z. Anat. Entwicklungsgesch. 139, 127–161 (1973).

42. Seltzer, B. & Pandya, D. N. Afferent cortical connections and architectonics of the superior temporal sulcus and surrounding cortex in the rhesus monkey. Brain Res. 149, 1–24 (1978).

43. Rosene, D. L. & Pandya, D. N. Architectonics and connections of the posterior parahippocampal gyrus in the rhesus monkey. Soc Neurosci Abstr 9:222, (1983).

44. Cherven, K. Mastering Gephi Network Visualization. (Pakt Publishing, 2015).

45. Zalesky, A., Fornito, A. & Bullmore, E. T. Fundamentals of Brain Network Analysis. Fundamentals of Brain Network Analysis (2016). doi:10.1016/C2012-0-06036-X

46. Barabási, A. & Albert, R. Emergence of Scaling in Random Networks. Science (80-.). 286, 509–512 (1999).

47. Eguíluz, V. M., Chialvo, D. R., Cecchi, G. A., Baliki, M. & Apkarian, A. V. Scale-free brain functional networks. Phys. Rev. Lett. 94, (2005).

48. Varshney, L. R., Chen, B. L., Paniagua, E., Hall, D. H. & Chklovskii, D. B. Structural properties of the Caenorhabditis elegans neuronal network. PLoS Comput. Biol. 7, (2011).

49. van den Heuvel, M. P., Stam, C. J., Boersma, M. & Hulshoff Pol, H. E. Small-world and scale-free organization of voxel-based resting-state functional connectivity in the human brain. Neuroimage 43, 528–539 (2008).

50. Sporns, O. & Zwi, J. D. The small world of the cerebral cortex. Neuroinformatics 2, 145–162 (2004).

51. Modha, D. S. & Singh, R. Network architecture of the long-distance pathways in the macaque brain. Proc. Natl. Acad. Sci. 107, 13485–13490 (2010).

52. Humphries, M.., Gurney, K. & Prescott, T.. The brainstem reticular formation is a small-world, not scale-free, network. Proc. R. Soc. B Biol. Sci. 273, 503–511 (2006).

53. Clauset, A., Shalizi, C. R. & Newman, M. E. J. Power-Law Distributions in Empirical Data. SIAM Rev. 51, 661–703 (2009).

54. Amaral, L. A. N. et al. Classes of small-world networks. Proc. Natl. Acad. Sci. U. S. A. 97, 11149–52 (2000).

55. Alstott, J., Bullmore, E. & Plenz, D. Powerlaw: A python package for analysis of heavy-tailed distributions. PLoS One 9, (2014).

56. Jacob, S. N. & Daniel, H. Structuring of Abstract Working Memory Content by Fronto-parietal Synchrony in Primate Cortex Article Structuring of Abstract Working Memory Content by Fronto-parietal Synchrony in Primate Cortex. 588–597 (2018). doi:10.1016/j.neuron.2018.07.025

57. Milo, R. Network Motifs: Simple Building Blocks of Complex Networks. Science (80-.). 298, 824–827 (2002).

58. Sporns, O. & Kötter, R. Motifs in Brain Networks. PLoS Biol. 2, (2004).

59. Harriger, L., van den Heuvel, M. P. & Sporns, O. Rich Club Organization of Macaque Cerebral Cortex and Its Role in Network Communication. PLoS One 7, (2012).

60. Honey, C. J., Kötter, R., Breakspear, M. & Sporns, O. Network structure of cerebral cortex shapes functional connectivity on multiple time scales. Proc. Natl. Acad. Sci. 104, 10240–10245 (2007).

61. Gollo, L. L., Mirasso, C. R., Atienza, M., Crespo-Garcia, M. & Cantero, J. L. Theta band zero-lag long-range cortical synchronization via hippocampal dynamical relaying. PLoS One 6, (2011).

62. Watts, D. J. & Strogatz, S. H. Collective dynamics of ‘small-world’ networks. Nature 393, 440–442 (1998).

63. Rubinov, M., Ypma, R. J. F., Watson, C., Bullmore, E. T. & Raichle, M. E. Wiring cost and topological participation of the mouse brain connectome. Proc. Natl. Acad. Sci. U. S. A. 112, 10032–10037 (2015).

64. Asmussen, S. Applied probability and queues. Second. (Springer-Verlag, 2003).

65. Mathworks®. Curve Fitting Toolbox™: User’s Guide (R2019b). MATLAB Manual (2019).

66. Moreno, Y., Nekovee, M. & Vespignani, A. Efficiency and reliability of epidemic data dissemination in complex networks. Phys. Rev. E - Stat. Physics, Plasmas, Fluids, Relat. Interdiscip. Top. 69, 4 (2004).

67. Albert, R., Jeong, H. & Barabási, A. L. Error and attack tolerance of complex networks. Nature 406, 378–382 (2000).

68. Stokes, M. G., Buschman, T. J. & Miller, E. K. Dynamic Coding for Flexible Cognitive Control. 221–241 (2017).

69. Bassett, D. S. & Bullmore, E. T. Small-World Brain Networks Revisited. Neuroscientist 23, 499–516 (2017).

70. Watson, C. E. & Chatterjee, A. A bilateral frontoparietal network underlies visuospatial analogical reasoning. Neuroimage 59, 2831–2838 (2012).

71. Lago-Fernández, L. F., Huerta, R., Corbacho, F. & Sigüenza, J. A. Fast response and temporal coherent oscillations in small-world networks. Phys. Rev. Lett. 84, 2758–2761 (2000).

72. Masuda, N. & Aihara, K. Global and local synchrony of coupled neurons in small-world networks. Biol. Cybern. 90, 302–309 (2004).

73. Bassett, D. S. & Sporns, O. Network neuroscience. Nat. Neurosci. 20, 353–364 (2017).

74. Schuman, C. D. et al. A Survey of Neuromorphic Computing and Neural Networks in Hardware. 1–88 (2017).

75. Bonin, G. Von & Bailey, P. The neocortex of Macaca Mulatta. University of Illinois Press (1947).

76. Brodmann, K. Vergleichende Lokalisationslehre der Grosshirnrinde in ihren Prinzipien dargestellt auf Grund des Zellenbaues. (1909).

77. Economo, C. von & Koskinas, G. Die cytoarchitektonik der hirnrinde des erwachsenen menschen. (1925).

78. Felleman, D. J. & Van Essen, D. C. Distributed hierarchical processing in the primate cerebral cortex. Cereb. Cortex 1, 1–47 (1991).

79. Pandya, D. N. & Yeterian, E. H. Architecture and Connections of Cortical Association Areas. in 3–61 (1985). doi:10.1007/978-1-4757-9619-3_1

80. Vogt, C. & Vogt, O. Allgemeine Ergebnisse unserer Hirnforschung. J. Psychol. Neurol. (1919).

81. Walker, A. E. A cytoarchitectural study of the prefrontal area of the macaque monkey. J. Comp. Neurol. 73, 59–86 (1940).

82. Goldman-Rakic, P. S. Circuitry of Primate Prefrontal Cortex and Regulation of Behavior by Representational Memory. in Comprehensive Physiology (John Wiley & Sons, Inc., 2011). doi:10.1002/cphy.cp010509

83. Petrides, M. Lateral prefrontal cortex: architectonic and functional organization. Philos. Trans. R. Soc. B Biol. Sci. 360, 781–795 (2005).

84. Bullmore, E. & Sporns, O. Complex brain networks: Graph theoretical analysis of structural and functional systems. Nature Reviews Neuroscience (2009). doi:10.1038/nrn2575

85. Lavail, J. H. & Lavail, M. M. Retrograde Axonal Transport in the Central Nervous System. Science (80-.). 176, 1416–1417 (1972).

86. Cowan, W. M., Gottlieb, D. I., Hendrickson, A. E., Price, J. L. & Woolsey, T. A. The autoradiographic demonstration of axonal connections in the central nervous system. Brain Res. 37, 21–51 (1972).

87. Cawthon, L. K. Primate Factsheets: Long-tailed macaque (Macaca fascicularis) Taxonomy, Morphology, & Ecology. Primate Info Net (2006). Available at: http://pin.primate.wisc.edu/factsheets/entry/long-tailed_macaque. (Accessed: 3rd November 2019)

88. Cawthon, L. K. Primate Factsheets: Rhesus macaque (Macaca mulatta) Taxonomy, Morphology, & Ecology. Primate Info Net (2005). Available at: http://pin.primate.wisc.edu/factsheets/entry/rhesus_macaque. (Accessed: 3rd November 2019)

89. Cawthon, L. K. Primate Factsheets: Pigtail macaque (Macaca nemestrina) Taxonomy, Morphology, & Ecology. Primate Info Net (2005). Available at: http://pin.primate.wisc.edu/factsheets/entry/pigtail_macaque/taxon. (Accessed: 3rd November 2019)

90. Cawthon, L. K. Primate Factsheets: Squirrel monkey (Saimiri) Taxonomy, Morphology, & Ecology. Primate Info Net (2006). Available at: http://pin.primate.wisc.edu/factsheets/entry/squirrel_monkey. (Accessed: 3rd November 2019)

91. Cawthon, L. K. Primate Factsheets: Owl monkey (Aotus) Taxonomy, Morphology, & Ecology. Primate Info Net (2005). Available at: http://pin.primate.wisc.edu/factsheets/entry/owl_monkey. (Accessed: 3rd November 2019)

92. Bakker, R., Wachtler, T. & Diesmann, M. CoCoMac 2.0 and the future of tract-tracing databases. Front. Neuroinform. 6, (2012).

93. Stephan, K. E. et al. Advanced database methodology for the Collation of Connectivity data on the Macaque brain (CoCoMac). Philos. Trans. R. Soc. London. Ser. B Biol. Sci. 356, 1159–1186 (2001).

94. Medalla, M. & Barbas, H. Diversity of laminar connections linking periarcuate and lateral intraparietal areas depends on cortical structure. Eur. J. Neurosci. 23, 161–179 (2006).

95. Bastian, M., Heymann, S. & Jacomy, M. Gephi: An Open Source Software for Exploring and Manipulating Networks. Third Int. AAAI Conf. Weblogs Soc. Media 361–362 (2009). doi:10.1136/qshc.2004.010033

96. Rubinov, M. & Sporns, O. Complex network measures of brain connectivity: Uses and interpretations. Neuroimage 52, 1059–1069 (2010).

97. Maslov, S. & Sneppen, K. Specificity and stability in topology of protein networks. Science (80-.). 296, 910–913 (2002).

98. Milo, R., Kashtan, N., Itzkovitz, S., Newman, M. E. J. & Alon, U. On the uniform generation of random graphs with prescribed degree sequences. (2003).

99. Dijkstra, E. W. A note on two problems in connexion with graphs. Numer. Math. 1, 269–271 (1959).

100. Telesford, Q. K., Joyce, K. E., Hayasaka, S., Burdette, J. H. & Laurienti, P. J. The Ubiquity of Small-World Networks. Brain Connect. 1, 367–375 (2011).

