## Supplementary Material for "Organization of Areal Connectivity in the Monkey Frontoparietal Network"

### Supplementary Discussion

The FPN's connection density is 45.86%. This qualifies it as a moderately dense network, with almost half of all possible interareal connections existing.

The binary directed adjacency matrix of the FPN provides both in and out-degree distributions which detail the number of incoming and outgoing connections to each area, respectively (Supplementary Figure 1). Each distribution has an average degree of 13.3. Most of the nodes have an in-degree similar in quantity to their out-degree. However, 8 of the nodes have a large difference between incoming and outgoing connections, potentially signaling functional specialization. The ratios of in-to-out-degree and out-to-in-degree were calculated to identify those areas exhibiting a disparity greater than 1.5x. Frontal area 6DR and parietal areas PGop, IPd and PFop have in-to-out-degree ratios of 1.73, 1.88, 1.83 and 2.67, respectively. Frontal area 46v and parietal areas PEc, PEa and PE have out-to-in-degree ratios of 1.53, 1.71, 2.5 and 3.5, respectively.

A Pareto chart is used to quantify the portion each factor contributes to an overall distribution. When applied to a total degree distribution, the chart can identify the portion each area contributes to overall connectivity. In this way, a Pareto chart of the total degree distribution provides a qualitative assessment of whether specific areas contribute more to the overall connectivity, thereby suggesting these areas may qualify as hubs (Supplementary Figure 2). In the FPN, the $80^{th}$ percentile of total connectivity is not reached until 20 of the 30 nodes of the network are accounted for, or 67.7% of the total network. In a network with hubs, the $80^{th}$ percentile would generally be reached much earlier in the chart, indicating that a small number of nodes account for a large percentage of the connectivity in the network. These are the nodes which would qualify as potential hubs. Therefore, there do not appear to be any hubs in the FPN according to this qualitative assessment.

Areas 45, 47/12 and PGm were found to participate in the M9 motif with a significantly greater frequency in the empirical FPN than in random ($p = 0$, 0 and 0, $z = 5.189$, 8.358 and 8.060) and lattice ($p = 0.046$, 0.014 and 0.029, $z = 1.878$, 2.440 and 2.109) networks (Supplementary Table 2). Interestingly, the structural connectivity profile of these areas form the M9 motif as well, with area 45 serving as the apex node and areas 47/12 and PGm serving as the outer nodes (see Figure 6 for M9 example).

**Areal Specialization**

Areas which have a great disparity between their total number of incoming and outgoing connections may have developed this topology through functional specialization. Specifically, nodes which have a high in-to-out-degree ratio suggest the node may be an information aggregator and distiller, acting as a filter for the nodes it sends projections to. Frontal area 6DR likely would be aggregating and disseminating information related to ocular motor movements to its target areas[1].

Nodes which have a high out-to-in-degree ratio suggest the node may be an information source for the nodes it sends projections to. Frontal area 46v has been reported to provide information to its target areas supporting a range of cognitive functions including visuospatial working memory[2].

The parietal areas identified may provide visual, visuospatial and sequential processing of somatosensory information in support of cognition such as visual and visuospatial working memory[2–6]. Parietal areas PGop, IPd and PFop may compile and filter the information while areas PEa, PEc and PE may act as information sources, thereby providing functional specialization in service of cognition. It is worth pointing out that 6 of the 8 areas which had a high difference between their incoming and outgoing connections were in the parietal region. This region may be under-explored as a result of not having enough tract-tracing studies with injections in the parietal areas of Pandya & Seltzer (1982). As a result, additional projections simply may not have been discovered yet.

**Connectivity**

There did not appear to be any areas servings as hubs in the FPN according to the Pareto chart. Hub areas would have produced a Pareto chart which reached the 80th percentile much earlier, while only a small percentage of the network was accounted for. For instance, the 50th percentile of the degree distribution of the incoming hyperlinks in the Web is reached after just 1.1% of the nodes in the network are accounted for. These nodes are considered hubs in the Web network (Newman, 2018). A more quantitative analysis was needed to properly classify the degree distributions of the FPN, which would yield insight into the likelihood of hubs[7,8].

**Motifs**

Synchrony in the FPN may originate from the over-represented frontal areas 45 and 47/12 and parietal area PGm. If all three areas were inactivated, it could be speculated that performance on cognitive tasks requiring synchrony in the FPN would be impaired. This would serve as an interesting follow-up experiment.

The M13 motif was very close to being significantly overrepresented. This motif differs from M9 through the addition of a reciprocal connection between its driven nodes. This serves to encourage non-zero lag synchrony and fails to promote zero lag synchrony due to frustration[9]. If the M13 motif had been significant, it would be interpreted as providing the FPN further topological flexibility to establish the kind of synchrony necessary for cognition.

### Small World

This work focused on an established neural network observed in isolation from the rest of the brain. When whole-brain connectomes are analyzed at the macroscale, they generally present with hub nodes and power-law scaling of their degree distributions[10,11]. This is likely because the heterogeneity of connectivity is much greater at the level of the whole brain. When analyzed in this way, there are a smaller number of highly connected nodes compared with a larger number of nodes with few connections.

### Supplementary Figures

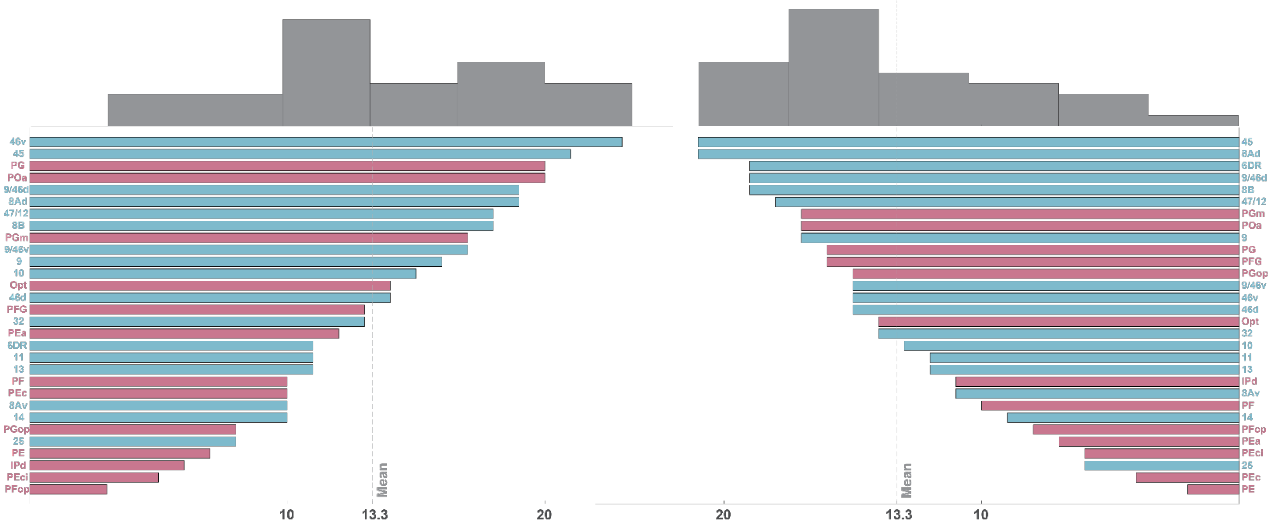


**Supplementary Figure 1**. The degree distributions for the frontoparietal network. Out-degree is on the left and in-degree is on the right. The average degree for both distributions is 13.3. Frontal areas are colored blue and parietal areas are pink.

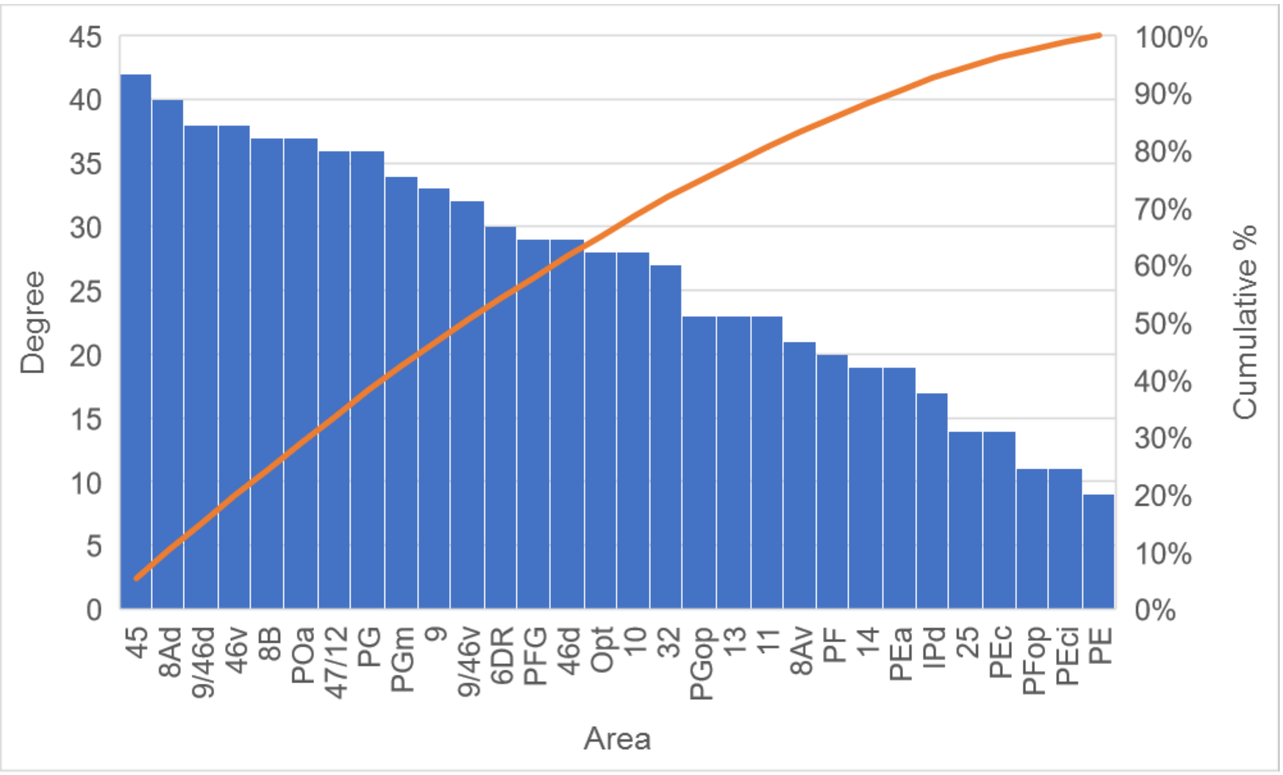


**Supplementary Figure 2.** A pareto chart of the total degree (in-degree + out-degree) for each node in the frontoparietal network. Totaling the degree counts for 20 of the 30 areas, up to area 11, accounts for 80% of overall connectivity in the network. This represents a traversal of about 67% of the network before reaching 80% total connectivity, which means it is unlikely that there are any hub nodes.

### Supplementary Tables

| | | Frontal | | | | | | | | | | | | | | | | |
|---|---|---|---|---|---|---|---|---|---|---|---|---|---|---|---|---|---|---|
| | | **10** | **9** | **32** | **14** | **25** | **8B** | **8Ad** | **9/46d** | **46d** | **46v** | **9/46v** | **8Av** | **45** | **47/12** | **13** | **11** | **6DR** |
| Frontal | **10** | | 12–15 | 12,13,15–17 | 12,13,15,16 | 15 | 13–15 | 15,17,18 | 14,18 | 12,14,15 | 15 | | | 15,19,20 | 12,15–17,20,21 | 15,16,21 | 12,15–17 | 15 |
| | **9** | 13,15,16 | | 12,13,15–17 | 13,15,16 | 15 | 13–15 | 12,14,15,22 | 14,15,18 | 14,15 | 15 | | | 19,20 | 15–17,20,21 | 15,21 | 15–17 | 13,15,23 |
| | **32** | 12,13,15,16,24 | 12–15,24 | | 12,13,15,16,24 | 12,15,24 | 13–15 | | 14 | 12,14,15,24 | 12,15 | | | 20 | 12,15–17,20,21,24 | 12,15,16,21,24 | 15–17,24 | 15 |
| | **14** | 13,16 | 13,14 | 12,13,16,17 | | | 13,14 | | 14 | 14 | | | | 20 | 16,17,20,21 | 16,21 | 16,17 | |
| | **25** | 13,16 | 13 | 12,13,16,17 | 13,16 | | 13 | | | | | | | | 21 | 16,21 | 17 | |
| | **8B** | 13 | 13,14,25 | 17,25 | | 13 | | 14,17,18,22,25 | 14,18 | 14,25,26 | 25 | 17 | 14 | 19,20 | 14,17,21,25 | 21 | 25 | 14,23,25,27,28 |
| | **8Ad** | 13,25 | 12–14,25 | 17 | | | 12,13,25,29 | | 12,14,18,25,29,30 | 12,14,25,26,29 | 12 | 12,17,29 | 12,14,25,29 | 12,20,25,29 | 12,17,25 | | | 12,23,25,27–32 |
| | **9/46d** | | 13,14,25 | | | | 14,25 | 14,17,18,22,25,30 | | 14,25 | 25 | 17 | 25 | 19,20,25 | 17,25 | 21 | | 14,23,25,27,28,32 |
| | **46d** | 13 | 13,14 | 12,13,17 | 13 | | 13,14 | 14,17,18,22 | 14,18 | | | | 14 | 19,20 | 17,20 | | | 23 |
| | **46v** | 13 | 12–14 | 13,17 | 13 | | 12–14 | 14,17,22 | 12,14 | 14,26 | | 17 | 12,14 | 12,19,20 | 12,17,20,21 | 12,21 | 17 | 12,28 |
| | **9/46v** | | | | | | 25 | 14,17,22 | 25 | | 25,26 | | 25,30 | 19,20,25 | 17,20,21,25 | 21,25 | 17,25 | 25 |
| | **8Av** | | | | | | 25 | 14,17,18,25 | | | | 17,30 | | 20,25,30 | 17,20,30 | | | 32 |
| | **45** | 19 | 19 | 17 | | | 14,19 | 14,17,18,22 | 14,19 | 14,19,26 | 19 | 17,19 | 14,30 | | 16,17,19,20 | 16,19 | 16,17,19 | 23,27,31,32 |
| | **47/12** | 12,13,16 | 12–14 | 12,13,16,17 | 12,13,16 | 12 | 12–14 | 14,17 | 14 | 12,14 | 12,26 | 17,20 | 14,30 | 12,19,20 | | 12,16,21 | 12,16,17 | |
| | **13** | 12,13,16 | 12–14 | 12,13,16,17 | 12,13,16 | 12 | 13,14 | | 14 | | | 20 | | 12,19,20 | 12,16,17,20,21 | | 12,16,17 | |
| | **11** | 13,15,16 | 13,14 | 12,13,16,17 | 13,15,16 | | | | 14 | 14 | 15,26 | 15,20 | | 15,19,20 | 15–17,20,21 | 15,16,21 | | |
| | **6DR** | | 13,14,23 | 17 | | | 13,14,23 | 12,14,17,18,22,23,30,31 | 12,14,23 | | | 17,33 | | 20,31 | | | | |
| Parietal | **PE** | | | | | | | 34 | | | | | | | | | | |
| | **PEci** | | | | | | | 17 | | | | | | | | | | 14 |
| | **PEc** | | | | | | | | 34 | | | | | | | | | 14,34 |
| | **PEa** | | | | | | | 14,22,34 | | | | | | | | | | 14,28 |
| | **PF** | | | | | | | | | | | 4,20,34,35 | 34,36 | 36 | 17,20,36 | | | |
| | **PFop** | | | | | | | | | | | | | | | | | |
| | **PFG** | | | | | | | 5,35 | | | 4 | 4,5,20,34,35 | | 4,5,19 | 4,5,17 | | | 5 |
| | **IPd** | | | | | | | 17,22 | | | | | | | | | | |
| | **POa** | | 34 | | | | 14,34 | 5,14,17,22,30,34,36–38 | 5,33,34,37,38 | 34 | 5,34,39 | 5,17,20,33,34,39 | 5,14,34,36,38,39 | 5,19,20,34 | 5,17,39 | | | 5 |

| | | Frontal | | | | | | | | | | | | | | | | |
|---|---|---|---|---|---|---|---|---|---|---|---|---|---|---|---|---|---|---|
| | | 10 | 9 | 32 | 14 | 25 | 8B | 8Ad | 9/46d | 46d | 46v | 9/46v | 8Av | 45 | 47/12 | 13 | 11 | 6DR |
| | PG | | | | | | 14,36 | 17,22,34–36,38,40 | 14,35,36,38 | 26,36 | 4,26,34 | 4,17,35,36 | 14,35,38 | 20,35,36 | | | 17,36 | 14,35 |
| | PGm | | 5,41 | 17 | | | 5,14,41 | 5,17,34,41 | 5,14,41 | 5,14,41 | | | | 5 | | | | 1,5,14,28,34 |
| | PGop | | | | | | | 14,34 | | | 34 | | | | | | | |
| | Opt | 5 | 5 | | | | 4,5,14 | 4,5,14 | 4 | 5,14,26 | 4,5,26 | 4 | | 4,5,19 | | | | 4 |

| | | Parietal | | | | | | | | | | | | |
|---|---|---|---|---|---|---|---|---|---|---|---|---|---|---|
| | | **PE** | **PEci** | **PEc** | **PEa** | **PF** | **PFop** | **PFG** | **IPd** | **POa** | **PG** | **PGm** | **PGop** | **Opt** |
| **Frontal** | **10** | | | | | | | | | | | | | 5 |
| | **9** | | | | | | | | | | | 5,41 | | 5 |
| | **32** | | | | | | | | | | | | | |
| | **14** | | | | | | | | | | | | | |
| | **25** | | | | | | | | | | | | | |
| | **8B** | | | | | | | | | | 36 | 5,25 | | 4,5 |
| | **8Ad** | | | | | | | 5 | 25,42 | 5,25,30,37,38,42 | 25,36,38,40,43,44 | 5,25,41 | 45 | 4,5,25 |
| | **9/46d** | | 25 | 25 | | | | | 25,42 | 5,25,37,38 | 25,36,38,40,44 | 5,25,41 | 25 | 4 |
| | **46d** | | | | | | | | | | 36,40,43,44 | 5,41 | | 5 |
| | **46v** | | | | | 43 | 45 | 4 | 42 | 5 | 4,40,43,44 | | 45 | 4,5 |
| | **9/46v** | | | | | 4,25,43 | | 4,5,25 | | 5,39 | 4,25,36,44 | 5 | 45 | 4 |
| | **8Av** | | | | | 36 | | | 25,42 | 5,25,38,39,42,43 | 38 | | | |
| | **45** | | | | | 36 | 45 | 4,5,19 | | 5,19 | 36,44 | 5 | | 4,5,19 |
| | **47/12** | | | | | 36 | | 4,5 | | 5,39 | | | | |
| | **13** | | | | | | | | | | | | | |
| | **11** | | | | | | | | | | | | | |
| | **6DR** | | | | | | | 5 | | 5 | | 5 | | 4,5 |
| **Parietal** | **PE** | | 3 | | 3,37,46 | | 3 | 6 | | | | 6 | 45 | |
| | **PEci** | | | | | | | 6 | | | 4 | 6 | | |
| | **PEc** | 3 | 3 | | 3,46 | | | 6 | 46 | | 3,4 | 3,6,41 | 3 | |
| | **PEa** | | 3 | 3 | | 4 | 45 | 4,6 | 42 | 6,42 | 4,36 | 6,41 | 3 | |
| | **PF** | | | | 4 | | 3,45 | 3,4,6 | 4 | 3,4,37,39,46 | | | 4 | |
| | **PFop** | | | | | | | 6 | | 6 | | | 45 | |
| | **PFG** | | | | 4,37 | 4 | 45 | | 4,42,46 | 4,6,39,46 | 4 | | 4,45 | |
| | **IPd** | | | | | 4 | | 4 | | 6,37,42 | | 6 | 45 | |
| | **POa** | | | | 6,39 | 4 | 6,45 | 4,6 | 37,42 | | 4,6,36,37,44 | 6 | 6,45 | 4,6 |
| | **PG** | | 4 | 4 | 4,36 | | 45 | 4,6 | 46 | 4,6,36,37,39 | | 1,6,36,41 | 4,45 | 4,6 |
| | **PGm** | 41 | 6 | 6 | 6,37,46 | | | | 6,46 | 6,37 | 1,6,36,41 | | 6 | 4,6 |
| | **PGop** | | | | | 4 | | 4,6 | | 6 | 4 | 6 | | 4 |
| | **Opt** | | | | | | | | | 4,6 | 4,6,44 | 4,6 | 4 | |

**Supplementary Table 1.** Association matrix for the frontoparietal network using the Petrides & Pandya (2007) parcellation scheme. The fields are populated with the references where a connection was identified. Empty fields signify the absence of any reports of connectivity. However, this does not imply that there is no connection between those areas.

| | Motif Class ID | | | | | | | | | | | | |
|---|---|---|---|---|---|---|---|---|---|---|---|---|---|
| | 1 | 2 | 3 | 4 | 5 | 6 | 7 | 8 | **9** | 10 | 11 | 12 | 13 |
| p-val rand | 1.000 | 1.000 | 1.000 | 1.000 | 1.000 | 1.000 | 1.000 | 1.000 | **0.000** | 1.000 | 1.000 | 1.000 | 0.000 |
| p-val latt | 0.528 | 0.659 | 0.029 | 0.994 | 0.016 | 1.000 | 0.073 | 1.000 | **0.049** | 0.710 | 0.683 | 0.794 | 0.054 |
| z-score rand | -8.269 | -10.687 | -8.087 | -5.993 | -5.830 | -6.106 | -5.375 | -8.149 | **12.032** | -10.516 | -7.755 | -7.863 | 19.122 |
| z-score latt | -0.155 | -0.516 | 1.864 | -2.504 | 2.280 | -4.169 | -0.267 | -3.401 | **1.612** | -0.660 | -0.589 | -0.825 | 1.611 |

**Supplementary Table 2.** Results from a permutation test comparing the frequency of occurrence of motif classes in the empirical FPN with their frequency in 100,000 random and lattice networks. Motif class ID 9 (bolded) was found to be significantly overrepresented in the empirical FPN in comparison to both random (p = 0, z = 12.032) and lattice null networks (p = 0.049, z = 1.612).

| | | Area | | | | | | | | | | | | | | | | | | | | | | | | | | | | | |
|---|---|---|---|---|---|---|---|---|---|---|---|---|---|---|---|---|---|---|---|---|---|---|---|---|---|---|---|---|---|---|---|
| | | 10 | 9 | 32 | 14 | 25 | 8B | 8Ad | 9/46d | 46d | 46v | 9/46v | 8Av | **45** | **47/12** | 13 | 11 | 6DR | PE | PEci | PEc | PEa | PF | PFop | PFG | IPd | POa | PG | **PGm** | PGop | Opt |
| Motif Class ID | 1 | 0.997 | 1.000 | 1.000 | 1.000 | 1.000 | 0.999 | 0.999 | 0.840 | 0.994 | 0.374 | 0.987 | 0.991 | 0.965 | 1.000 | 1.000 | 1.000 | 0.851 | 1.000 | 0.977 | 1.000 | 0.996 | 1.000 | 0.490 | 1.000 | 0.901 | 0.995 | 0.978 | 1.000 | 0.859 | 1.000 |
| | | *0.307* | *0.860* | *0.862* | *0.623* | *0.624* | *0.745* | *0.861* | *0.235* | *0.397* | *0.139* | *0.304* | *0.232* | *0.623* | *0.863* | *0.861* | *0.625* | *0.023* | *0.142* | *0.232* | *0.139* | *0.045* | *0.622* | *0.062* | *0.862* | *0.139* | *0.626* | *0.398* | *0.863* | *0.062* | *0.860* |
| | 2 | 1.000 | 1.000 | 1.000 | 1.000 | 1.000 | 1.000 | 0.997 | 0.659 | 1.000 | 1.000 | 1.000 | 1.000 | 0.998 | 1.000 | 1.000 | 1.000 | 1.000 | 1.000 | 1.000 | 1.000 | 1.000 | 1.000 | 1.000 | 1.000 | 1.000 | 1.000 | 1.000 | 1.000 | 1.000 | 1.000 |
| | | *0.409* | *0.905* | *0.122* | *0.172* | *0.409* | *0.783* | *0.312* | *0.001* | *0.409* | *0.907* | *0.787* | *0.170* | *0.526* | *0.907* | *0.905* | *0.086* | *0.524* | *0.236* | *0.124* | *0.523* | *0.526* | *0.906* | *0.522* | *0.786* | *0.171* | *0.787* | *0.906* | *0.788* | *0.232* | *0.905* |
| | 3 | 1.000 | 1.000 | 1.000 | 0.996 | 1.000 | 1.000 | 0.966 | 0.930 | 0.997 | 0.608 | 1.000 | 0.998 | 0.997 | 1.000 | 0.997 | 1.000 | 0.485 | 0.797 | 0.974 | 0.916 | 0.924 | 0.999 | 0.995 | 1.000 | 0.973 | 1.000 | 0.945 | 1.000 | 1.000 | 1.000 |
| | | *0.845* | *0.844* | *0.844* | *0.227* | *0.845* | *0.844* | *0.550* | *0.310* | *0.308* | *0.001* | *0.845* | *0.161* | *0.844* | *0.846* | *0.163* | *0.420* | *0.007* | *0.227* | *0.046* | *0.165* | *0.086* | *0.312* | *0.007* | *0.845* | *0.004* | *0.846* | *0.165* | *0.845* | *0.119* | *0.846* |
| | 4 | 0.988 | 0.999 | 0.855 | 0.602 | 1.000 | 1.000 | 0.068 | 0.784 | 0.994 | 0.908 | 0.875 | 0.994 | 0.999 | 1.000 | 1.000 | 0.936 | 0.171 | 0.889 | 0.161 | 0.994 | 0.974 | 0.998 | 0.989 | 0.543 | 0.938 | 0.566 | 0.973 | 0.886 | 0.845 | 0.984 |
| | | *0.823* | *0.847* | *0.546* | *0.329* | *0.953* | *0.825* | *0.005* | *0.298* | *0.848* | *0.848* | *0.551* | *0.870* | *0.734* | *0.963* | *0.977* | *0.702* | *0.080* | *0.216* | *0.330* | *0.394* | *0.547* | *0.887* | *0.978* | *0.268* | *0.866* | *0.328* | *0.770* | *0.472* | *0.733* | *0.798* |
| | 5 | 1.000 | 1.000 | 1.000 | 0.999 | 0.999 | 0.999 | 0.981 | 0.992 | 0.994 | 1.000 | 1.000 | 0.972 | 0.998 | 0.999 | 1.000 | 1.000 | 1.000 | 0.354 | 0.547 | 0.975 | 0.959 | 0.998 | 0.956 | 0.996 | 0.986 | 1.000 | 1.000 | 1.000 | 0.996 | 1.000 |
| | | *0.626* | *0.624* | *0.627* | *0.626* | *0.628* | *0.626* | *0.131* | *0.232* | *0.132* | *0.038* | *0.624* | *0.038* | *0.629* | *0.627* | *0.626* | *0.629* | *0.038* | *0.001* | *0.004* | *0.019* | *0.004* | *0.388* | *0.070* | *0.134* | *0.038* | *0.629* | *0.630* | *0.627* | *0.009* | *0.625* |
| | 6 | 0.999 | 0.986 | 0.908 | 1.000 | 1.000 | 0.972 | 0.993 | 0.329 | 0.851 | 0.136 | 0.815 | 0.573 | 0.990 | 1.000 | 0.998 | 0.874 | 0.941 | 0.966 | 0.998 | 0.978 | 0.996 | 0.974 | 0.395 | 0.997 | 0.849 | 0.942 | 0.803 | 0.867 | 0.931 | 0.990 |
| | | *0.918* | *0.798* | *0.666* | *0.939* | *0.992* | *0.724* | *0.821* | *0.218* | *0.601* | *0.038* | *0.510* | *0.384* | *0.722* | *0.891* | *0.875* | *0.606* | *0.800* | *0.979* | *0.841* | *0.941* | *0.918* | *0.753* | *0.106* | *0.892* | *0.354* | *0.511* | *0.355* | *0.573* | *0.603* | *0.840* |
| | 7 | 0.994 | 0.994 | 0.995 | 0.991 | 0.974 | 0.990 | 0.981 | 0.988 | 0.995 | 0.936 | 0.993 | 0.993 | 0.975 | 0.991 | 0.993 | 0.994 | 0.973 | 0.776 | 0.951 | 0.929 | 0.981 | 0.992 | 0.889 | 0.994 | 0.975 | 0.986 | 0.985 | 0.993 | 0.978 | 0.995 |
| | | *0.007* | *0.007* | *0.008* | *0.008* | *0.008* | *0.008* | *0.008* | *0.007* | *0.008* | *0.007* | *0.008* | *0.008* | *0.007* | *0.008* | *0.008* | *0.008* | *0.008* | *0.008* | *0.008* | *0.008* | *0.008* | *0.008* | *0.007* | *0.008* | *0.008* | *0.008* | *0.007* | *0.008* | *0.008* | *0.008* |
| | 8 | 0.999 | 1.000 | 1.000 | 0.771 | 0.981 | 1.000 | 0.999 | 0.999 | 0.998 | 0.980 | 1.000 | 0.999 | 1.000 | 1.000 | 0.999 | 0.998 | 1.000 | 0.690 | 0.378 | 0.987 | 0.975 | 0.990 | 0.535 | 0.938 | 0.841 | 0.997 | 1.000 | 1.000 | 0.969 | 1.000 |
| | | *0.791* | *0.705* | *0.890* | *0.327* | *0.790* | *0.504* | *0.253* | *0.505* | *0.506* | *0.506* | *0.792* | *0.791* | *0.702* | *0.891* | *0.790* | *0.707* | *0.408* | *0.411* | *0.611* | *0.611* | *0.502* | *0.702* | *0.790* | *0.137* | *0.607* | *0.608* | *0.891* | *0.793* | *0.409* | *0.889* |
| | **9** | 0.000 | 0.000 | 0.009 | 0.279 | 0.001 | 0.001 | 0.000 | 0.021 | 0.001 | 0.004 | 0.000 | 0.000 | **0.000** | **0.000** | 0.000 | 0.021 | 0.000 | 0.132 | 0.130 | 0.003 | 0.007 | 0.000 | 0.141 | 0.000 | 0.005 | 0.000 | 0.000 | **0.000** | 0.000 | 0.000 |
| | | *0.379* | *0.198* | *0.596* | *0.936* | *0.594* | *0.241* | *0.125* | *0.465* | *0.396* | *0.464* | *0.092* | *0.382* | ***0.046*** | ***0.014*** | *0.318* | *0.681* | *0.431* | *0.995* | *0.968* | *0.783* | *0.700* | *0.232* | *0.987* | *0.062* | *0.716* | *0.188* | *0.078* | ***0.029*** | *0.448* | *0.097* |
| | 10 | 1.000 | 1.000 | 1.000 | 0.998 | 1.000 | 1.000 | 1.000 | 1.000 | 1.000 | 1.000 | 1.000 | 1.000 | 1.000 | 1.000 | 1.000 | 1.000 | 1.000 | 0.917 | 0.966 | 0.998 | 1.000 | 1.000 | 0.902 | 1.000 | 1.000 | 1.000 | 1.000 | 1.000 | 1.000 | 1.000 |
| | | *0.770* | *0.769* | *0.450* | *0.160* | *0.607* | *0.162* | *0.449* | *0.048* | *0.233* | *0.160* | *0.769* | *0.452* | *0.608* | *0.770* | *0.769* | *0.162* | *0.328* | *0.607* | *0.327* | *0.607* | *0.452* | *0.608* | *0.326* | *0.606* | *0.446* | *0.447* | *0.232* | *0.605* | *0.159* | *0.767* |
| | 11 | 0.996 | 1.000 | 0.994 | 0.973 | 0.964 | 1.000 | 1.000 | 0.814 | 0.990 | 1.000 | 1.000 | 0.969 | 0.999 | 1.000 | 1.000 | 0.932 | 0.937 | 0.620 | 0.668 | 0.002 | 0.408 | 0.995 | 0.748 | 1.000 | 0.818 | 1.000 | 0.929 | 1.000 | 0.998 | 1.000 |
| | | *0.340* | *0.852* | *0.433* | *0.539* | *0.746* | *0.647* | *0.745* | *0.015* | *0.339* | *0.103* | *0.743* | *0.432* | *0.540* | *0.853* | *0.852* | *0.267* | *0.105* | *0.854* | *0.540* | *0.104* | *0.147* | *0.649* | *0.261* | *0.854* | *0.147* | *0.749* | *0.010* | *0.744* | *0.539* | *0.853* |
| | 12 | 0.998 | 1.000 | 0.912 | 0.034 | 0.379 | 1.000 | 1.000 | 0.997 | 0.742 | 0.949 | 1.000 | 0.247 | 1.000 | 1.000 | 0.973 | 0.422 | 0.418 | 0.059 | 0.010 | 0.277 | 0.256 | 0.939 | 0.002 | 1.000 | 0.020 | 0.999 | 1.000 | 1.000 | 0.471 | 1.000 |
| | | *0.583* | *0.557* | *0.386* | *0.507* | *0.928* | *0.234* | *0.315* | *0.036* | *0.178* | *0.007* | *0.559* | *0.507* | *0.405* | *0.702* | *0.775* | *0.434* | *0.075* | *0.986* | *0.914* | *0.928* | *0.670* | *0.879* | *0.891* | *0.730* | *0.639* | *0.076* | *0.317* | *0.252* | *0.478* | *0.892* |
| | 13 | 0.000 | 0.000 | 0.000 | 0.000 | 0.000 | 0.000 | 0.000 | 0.000 | 0.000 | 0.000 | 0.000 | 0.000 | 0.000 | 0.000 | 0.000 | 0.000 | 0.000 | 0.047 | 0.055 | 0.001 | 0.010 | 0.000 | 0.018 | 0.000 | 0.007 | 0.000 | 0.000 | 0.000 | 0.000 | 0.000 |
| | | *0.334* | *0.082* | *0.449* | *0.765* | *0.787* | *0.027* | *0.009* | *0.140* | *0.422* | *0.160* | *0.189* | *0.720* | *0.000* | *0.035* | *0.526* | *0.643* | *0.527* | *0.996* | *0.962* | *0.923* | *0.869* | *0.752* | *0.961* | *0.514* | *0.873* | *0.127* | *0.226* | *0.371* | *0.748* | *0.188* |

**Supplementary Table 3.** Results from a permutation test comparing the frequency of participation of areas in motif classes in the empirical FPN with their frequency in 100,000 random and lattice networks. Random network p-values are represented in plain font and lattice network p-values are represented in italics. Bolded text highlights areas 45, 47/12 and PGm which were found to participate in motif class 9 with a significantly greater frequency in the FPN than in random (p = 0.0, z = 5.189, 8.358 and 8.060) and lattice networks (p = 0.046, 0.014 and 0.029, z = 1.878, 2.440 and 2.109).

**Supplementary References**


1. Luppino, G. & Rizzolatti, G. The Organization of the Frontal Motor Cortex. *News Physiol. Sci.* **15**, 219–224 (2000).

2. Goldman-Rakic, P. S. Circuitry of Primate Prefrontal Cortex and Regulation of Behavior by Representational Memory. in *Comprehensive Physiology* (John Wiley & Sons, Inc., 2011). doi:10.1002/cphy.cp010509

3. Pandya, D. N. & Seltzer, B. Intrinsic connections and architectonics of posterior parietal cortex in the rhesus monkey. *J. Comp. Neurol.* **204**, 196–210 (1982).

4. Rozzi, S. *et al.* Cortical connections of the inferior parietal cortical convexity of the macaque monkey. *Cereb. Cortex* **16**, 1389–1417 (2006).

5. Cavada, C. & Goldman-Rakic, P. S. Posterior parietal cortex in rhesus monkey: II. Evidence for segregated corticocortical networks linking sensory and limbic areas with the frontal lobe. *J. Comp. Neurol.* **287**, 422–445 (1989).

6. Cavada, C. & Goldman-Rakic, P. S. Posterior parietal cortex in rhesus monkey: I. Parcellation of areas based on distinctive limbic and sensory corticocortical connections. *J. Comp. Neurol.* **287**, 393–421 (1989).

7. Alstott, J., Bullmore, E. & Plenz, D. Powerlaw: A python package for analysis of heavy-tailed distributions. *PLoS One* **9**, (2014).

8. Clauset, A., Shalizi, C. R. & Newman, M. E. J. Power-Law Distributions in Empirical Data. *SIAM Rev.* **51**, 661–703 (2009).

9. Gollo, L. L., Mirasso, C., Sporns, O. & Breakspear, M. Mechanisms of Zero-Lag

Synchronization in Cortical Motifs. *PLoS Comput. Biol.* **10**, (2014).

10. Watts, D. J. & Strogatz, S. H. Collective dynamics of 'small-world' networks. *Nature* **393**, 440–442 (1998).

11. Oh, S. W. *et al.* A mesoscale connectome of the mouse brain. *Nature* **508**, 207–214 (2014).

12. Barbas, H. & Pandya, D. N. Architecture and intrinsic connections of the prefrontal cortex in the rhesus monkey. *J. Comp. Neurol.* **286**, 353–375 (1989).

13. Barbas, H., Ghashghaei, H. & Dombrowski, S. M. Medial Prefrontal Cortices Are Unified by Common Connections With Superior Temporal Cortices and Distinguished by Input From Memory-Related Areas in the Rhesus Monkey. **367**, 343–367 (1999).

14. Petrides, M. & Pandya, D. N. Dorsolateral prefrontal cortex: comparative cytoarchitectonic analysis in the human and the macaque brain and corticocortical connection patterns. *Eur. J. Neurosci.* **11**, 1011–1036 (1999).

15. Petrides, M. & Pandya, D. N. Efferent Association Pathways from the Rostral Prefrontal Cortex in the Macaque Monkey. *J. Neurosci.* **27**, 11573–11586 (2007).

16. Price, J. L., Carmichael, S. T. & Price, J. L. Connectional networks within the orbital and medial prefrontal cortex of macaque monkeys. *J. Comp. Neurol.* **371**, 179–207 (1996).

17. Barbas, H. Anatomic organization of basoventral and mediodorsal visual recipient prefrontal regions in the rhesus monkey. *J. Comp. Neurol.* **276**, 313–342 (1988).

18. Hackett, T. A., Stepniewska, I. & Kaas, J. H. Prefrontal connections of the parabelt auditory cortex in macaque monkeys. *Brain Res.* **817**, 45–58 (1999).

19. Gerbella, M. *et al.* Cortical connections of the macaque caudal ventrolateral prefrontal areas 45A and 45B. *Cereb. Cortex* **20**, 141–168 (2010).

20. Petrides, M. & Pandya, N. Comparative cytoarchitectonic analysis of human and macaque ventrolateral pfc and corticortical cnxn patterns in monkey. *Eur. J. Neurosci.* **16**, 291–310 (2001).

21. Barbas, H. Organization of cortical afferent input to orbitofrontal areas in the rhesus monkey. *Neuroscience* **56**, 841–864 (1993).

22. Barbas, H. & Mesulam, M. M. -M M. Organization of afferent input to subdivisions of area 8 in the rhesus monkey. *J. Comp. Neurol.* **200**, 407–431 (1981).

23. Barbas, H. & Pandya, D. N. Architecture and frontal cortical connections of the premotor cortex (area 6) in the rhesus monkey. *J. Comp. Neurol.* **256**, 211–228 (1987).

24. Pandya, D. N., Van Hoesen, G. W. & Mesulam, M. M. Efferent connections of the cingulate gyrus in the rhesus monkey. *Exp. Brain Res.* **42**, 319–330 (1981).

25. Petrides, M. & Pandya, D. N. N. Efferent association pathways originating in the caudal prefrontal cortex in the macaque monkey. *J. Comp. Neurol.* **498**, 227–251 (2006).

26. Jacobson, S. & Trojanowski, J. Q. Prefrontal granular cortex of the rhesus monkey. I. Intrahemispheric cortical afferents. *Brain Res.* **132**, 209–233 (1977).

27. Luppino, G. *et al.* Prefrontal and agranular cingulate projections to the dorsal premotor areas F2 and F7 in the macaque monkey. *Eur. J. Neurosci.* **17**, 559–578 (2003).

28. Ghosh, S. & Gattera, R. A comparison of the ipsilateral cortical projections to the dorsal and ventral subdivisions of the macaque premotor cortex. *Somatosens. Mot. Res.* **12**, 359–

378 (1995).

29. Stanton, G. B., Bruce, C. J. & Goldberg, M. E. Topography of projections to the frontal lobe from the macaque frontal eye fields. *J. Comp. Neurol.* **330**, 286–301 (1993).

30. Huerta, M. F., Krubitzer, L. A. & Kaas, J. H. O. N. H. Frontal eye field as defined by intracortical microstimulation in squirrel monkeys, owl monkeys, and macaque monkeys II. cortical connections. *J. Comp. Neurol.* **265**, 332–361 (1987).

31. Arikuni, T. O. M., Watanabe, K. & Kubota, K. Connections of area 8 with area 6 in the brain of the macaque monkey. *J. Comp. Neurol.* **277**, 21–40 (1988).

32. Luppino, G., Matelli, M. & Rizzolatti, G. Cortico-cortical connections of two electrophysiologically identified arm representations in the mesial agranular frontal cortex. *Exp. Brain Res.* **82**, 214–218 (1990).

33. Barbas, H. & Mesulam, M.-M. M. Cortical afferent input to the principals region of the rhesus monkey. *Neuroscience* **15**, 619–637 (1985).

34. Petrides, M. & Pandya, D. N. Projections to the frontal cortex from the posterior parietal region in the rhesus monkey. *J. Comp. Neurol.* **228**, 105–116 (1984).

35. Petrides, M. & Pandya, D. N. Distinct parietal and temporal pathways to the homologues of Broca’s area in the monkey. *PLoS Biol.* **7**, (2009).

36. Andersen, R. A., Asanuma, C., Essick, G. & Siegel, R. M. Corticocortical connections of anatomically and physiologically defined subdivisions within the inferior parietal lobule. *J. Comp. Neurol.* **296**, 65–113 (1990).

37. Blatt, G. J., Andersen, R. A. & Stoner, G. R. Visual receptive field organization and

cortico-cortical connections of the lateral intraparietal area (area LIP) in the macaque. *J. Comp. Neurol.* **299**, 421–445 (1990).

38. Medalla, M. & Barbas, H. Diversity of laminar connections linking periarcuate and lateral intraparietal areas depends on cortical structure. *Eur. J. Neurosci.* **23**, 161–179 (2006).

39. Borra, E. *et al.* Cortical Connections of the Macaque Anterior Intraparietal (AIP) Area. *Cereb. Cortex* **18**, 1094–1111 (2008).

40. Maioli, M. G., Squatrito, S., Samolsky-dekel, B. G. & Riva Sanseverino, E. Corticocortical connections between frontal periarcuate regions and visual areas of the superior temporal sulcus and the adjoining inferior parietal lobule in the macaque monkey. *Brain Res.* **789**, 118–125 (1998).

41. Parvizi, J. *et al.* Neural connections of the posteromedial cortex in the macaque. *Proc. Natl. Acad. Sci.* **103**, 1563–1568 (2006).

42. Lewis, J. W. & Van Essen, D. C. Corticocortical connections of visual, sensorimotor, and multimodal processing areas in the parietal lobe of the macaque monkey. *J. Comp. Neurol.* **428**, 112–137 (2000).

43. Neal, J. W., Pearson, R. C. A. & Powell, T. P. S. The ipsilateral corticocortical connections of area 7 with the frontal lobe in the monkey. *Brain Res.* **509**, 31–40 (1990).

44. Andersen, R. A., Asanuma, C. & Cowan, W. M. Callosal and prefrontal associational projecting cell populations in area 7A of the macaque monkey: A study using retrogradely transported fluorescent dyes. *J. Comp. Neurol.* **232**, 443–455 (1985).

45. Cipolloni, P. B. & Pandya, D. N. Cortical connections of the frontoparietal opercular areas

in the Rhesus monkey. *J. Comp. Neurol.* **403**, 431–458 (1999).

46. Seltzer, B. & Pandya, D. N. Posterior parietal projections to the intraparietal sulcus of the rhesus monkey. *Exp. Brain Res.* **62**, 459–469 (1986).
